# MePHD1.2 affects the synthesis of cyanogenic glycosides by regulating the transcription of *MeCYP79D2* in cassava

**DOI:** 10.1101/2024.04.18.590026

**Authors:** Mengtao Li, Xiao Zhao, Yuanchao Li, Xiaoye Zhao, Weitao Mai, Yajun Li, Qibing Liang, Qingchun Yin, Wenquan Wang, Jinping Liu, Xin Chen

## Abstract

The high content of cyanogenic glycosides (CG) in cassava storage tubers seriously affects human food safety. CG play crucial roles in plant growth and development and can protect cassava leaves from being masticated by herbivorous predators.

Nevertheless, the regulatory mechanism of CG biosynthesis, which results in a low CG content in storage tubers and high CG content in leaves, remains poorly understood.

Here, yeast one-hybrid assay was performed using a mixed cDNA library of cassava storage roots and leaves as prey and the promoter of *MeCYP79D2* as bait. MeCYP79D2, a cytochrome P450 protein, is the rate-limiting enzyme for CG synthesis in cassava. From this information, a candidate regulator of *MeCYP79D2*, that is, transcription factor MePHD1.2, was selected.

MePHD1.2, which is located in the nucleus and exhibits a transcription inhibitory activity, can directly bind to PD2 segment in the promoter of *MeCYP79D2*, which results in its repressed expression. In cassava, the transcriptional activity of *MeCYP79D2* was considerably enhanced in *mephd1.2* lines, which caused an increase in the contents of linamarin and lotaustralin.

Our findings unveil a novel regulatory module governing CG biosynthesis, wherein mutation of *MePHD1.2* attenuates its transcription inhibition on *MeCYP79D2* and boosts CGs biosynthesis in cassava.

## Introduction

Cassava (*Manihot esculenta* Crantz) belongs to the three major tuberous crops in the world, and it is widely cultivated in tropical regions. Its root is considered a staple food for nearly one billion people living in tropical and subtropical regions (Panghal *et al*., 2019). Cassava, whose leaves contain over 20% crude protein, exhibits an enormous potential for applications in animal feed and human food production. However, its aerial parts features high concentrations of cyanogenic glycosides (CG), which limit its utilisation in animal feed or human consumption (Latif & Müller, 2015).

CGs comprise plant products or secondary metabolites derived from amino acids (AAs) with oximes and cyanohydrins (α-hydroxynitriles) as key intermediates; these compounds are widely distributed in all tissues of cassava, mainly synthesised in the leaves and transported to the roots (Møller, 2010; Kannangara *et al*., 2011; Gleadow and Møller, 2014). CGs can help in the plant defence against external encroachment and serve as deterrents to herbivores (Zagrobelnya *et al*., 2004; Ballhorn *et al*., 2016). CGs, specifically linamarin and lotaustratalin, exist in all tissues of cassava. When hydrolysed, these compounds can release hydrogen cyanide (HCN) (Santana *et al*., 2002; Ogbonna *et al*., 2021). In cassava, CGs also function as storage compounds of reduced nitrogen and carbon and quenchers of reactive oxygen species (Frederik *et al*., 2018). However, long-term consumption of cassava storage roots with high CG contents can cause nausea, diarrhoea and other symptoms (Banea-Mayambu *et al*., 1997; Lebot, 2019). Therefore, cultivation of new cassava varieties with high CG amounts in leaves and low CG levels in storage roots is one of the currently important breeding goals.

Cytochrome P450s are characteristic enzymes that are ubiquitously present in numerous plants and animal species; they play important roles in various biochemical pathways (Zagrobelny *et al*., 2004). Most cytochrome P450s are associated with the biosynthesis of terpenoids, volatile derived fatty acids, plant hormones (gibberellins) and brassinosteroids (Dudereva *et al*., 2004; Fischbach and Clardy, 2007). In spite of species differences, CYP79s are the primary key enzymes in CG production, and they are considered rate-limiting enzymes of cyanogenesis pathways (Gleadow and Møller, 2014). The synthesis process of CGs involves three steps in cassava (Schmidt *et al*., 2018). Firstly, the dehydration and decarboxylation of L-valine and L-isoleucine occur via the catalysis of MeCYP79D1 and MeCYP79D2, which results in the production of (Z)-acetaldoxime, respectively. Subsequently, MeCYP71E7 catalyses (Z)-acetaldoxime to produce cyanohydrin, acetone cyanohydrin and 2-hydroxy-2-methylnitrile (Jørgensen *et al*., 2011). Finally, UDP glucosyltransferases, namely, MeUGT85K4 and MeUGT85K5, catalyse the production of linamarin and lotaustralin, respectively, from acetone cyanol and 2-hydroxy-2-methylnitrile (Fig. 1; Andersen *et al*., 2000; Kannangara *et al*., 2011). Hydrolysis of CGs occur via the continuous enzymatic reaction of β-glucosidase and α-hydroxynitrilase, which forms glucose, aldehydes or ketones and toxic hydrogen cyanide when cassava organs are suffering from mechanical damage (McMahon *et al*., 2022).

**Fig. 1.**
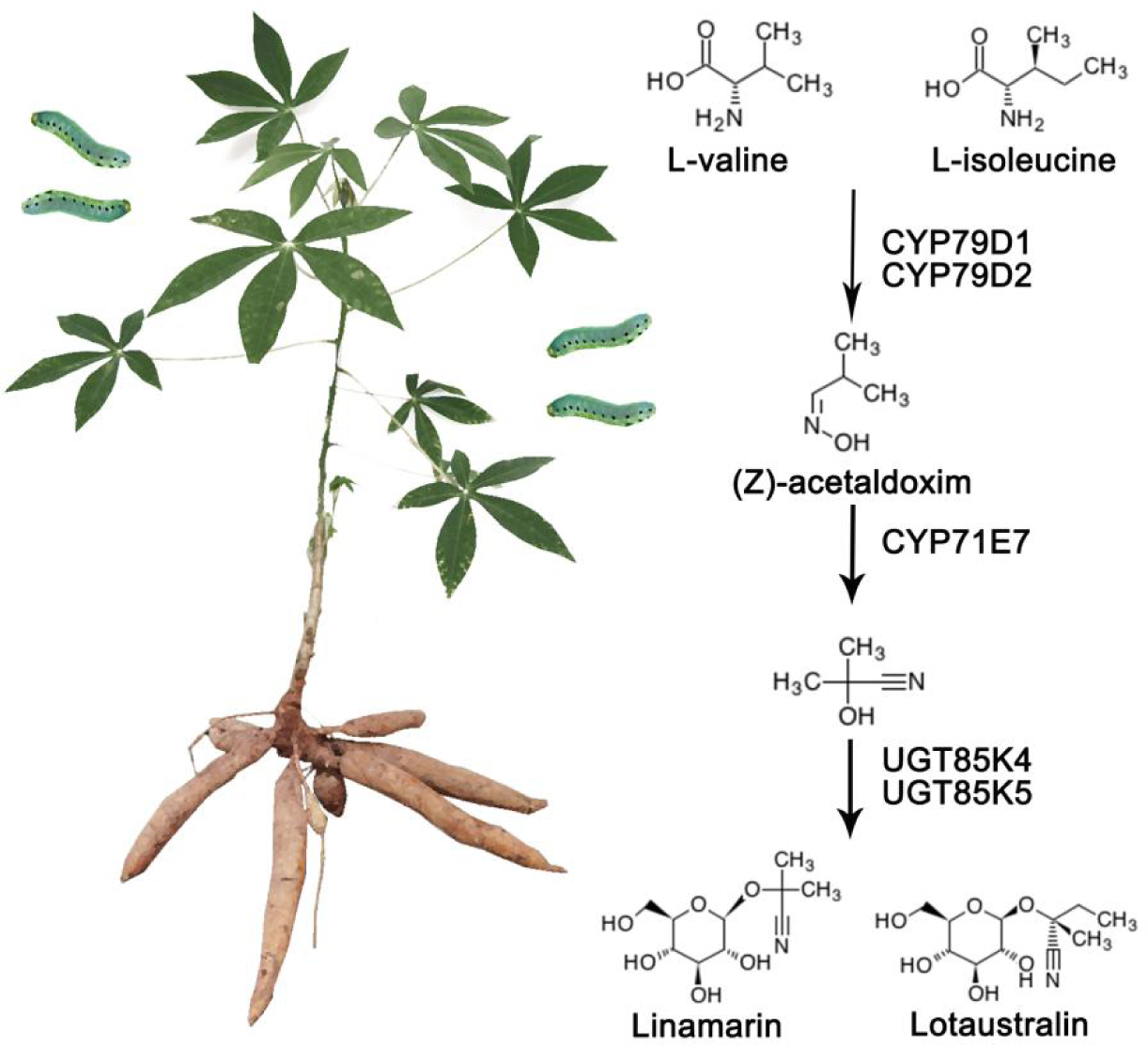
CG biosynthesis pathway in cassava.

When animal gnawing damage the plants of cassava, lima bean, sorghum and other crops, the expression levels of *CYP79Ds* genes are substantially up-regulated, and the synthesis efficiency of CGs and dhurrin increase, which reduces palatability of the plants and prevents further gnawing by animals (Lai *et al*., 2020; Easson *et al*., 2021). The simultaneous inhibition of the expressions of *MeCYP79D1* and *MeCYP79D2* through RNAi can greatly reduce CGs contents in cassava leaves and roots (Jørgensen *et al*., 2005). In addition, during the knockout of *MeCYP79D1* and *MeCYP79D2* by gene editing, the dual knockout lines showed no cyanogenic potential, and *MeCYP79D2* knockouts revealed a more drastic reduction in cyanogenic potential than *MeCYP79D1* knockouts and wild types, which indicates that *MeCYP79D2* is a dominant contributor to the biosynthesis of CGs in cassava (Gomez *et al*., 2023).

The transcription factor LjbHLH can up-regulate the transcriptional activity of *LjCYP79D3* in *Lotus japonicus* through induction of jasmonic acid and promote the synthesis of lotaustralin (Chen *et al*., 2022). In cucumber, the transcription factor CsBl causes the up-regulation of the expression of *CsBi* to promote cucurbitacine synthesis in leaves. Moreover, the alteration in Cys393Tyr in CsBl results in the reduced synthesis efficiency of cucurbitacine because CsBl cannot bind to the promoter of *CsBi*, which also *CsBi* encodes the enzyme responsible for the catalysis of the first and rate-limiting step in cucurbitacine synthesis pathway (Shang *et al*., 2014). Similarly, the alteration in Leu346Phe caused PdbHLH2 to form an unstable homodimer, as a result, its binding capability to the promoters of *PdCYP79D16* and *PdCYP79AN24* was reduced, the transcription activities of its targets were down-regulated, and the synthesis of amygdalin (a type of CG) in the endosperm of almonds, which led to the generation of sweet almonds, decreased (Sánchez-Pérez *et al*., 2019).

Plant homeodomain (PHD) refers to a class of zinc finger proteins that contain Cys4-His-Cys3 zinc-binding motifs and is involved in various molecular functions (Ma *et al*., 2018). The PHD-finger domain can specifically recognise the methylated modification of histone and may serve as an important reader of histone cryptogram (Jain *et al*., 2020). Early bolting in short days (EBS) of *Arabidopsis* involves two Bromo adjacent homology (BAH)-PHD domains that bind to H3K27me2 and H3K4me3 as a bivalent histone reader to mediate the switching of binding preferences between H3K27me2 and H3K4me3 and modulate flowering time in *Arabidopsis* (Qian *et al*., 2021). Alfin1-like proteins (ALs), which are members of the PHD-finger family, can read the epigenetic modification of histone, e.g. methylate H3K4me2/me3, repress the transcription activities of *AtbZIP50*, *AtCAX1*, *AtFAO* and *AtWRKY11* genes and intensify the tolerances to drought and salt. ALs commonly bind to the G-rich motif (GTGGAG), and the mutations of its AAs at positions 34 and 35 in AL6 caused the loss of ability to bind the G-rich motif (Wei *et al*., 2015).

However, available information on the PHD transcription factor involved in the synthesis of CG is limited. In the present study, the promoter of *MeCYP79D2* was cloned to construct a bait vector for yeast one-hybrid assay using a mixed cDNA library as prey, and the transcription factor MePHD1.2 was screened out. MePHD1.2 is a nucleus-localised protein, and it can bind to the PD2 segment in the *MeCYP79D2* promoter, down-regulate its transcriptional activity and increase the CG content in the leaves of *mephd1.2* mutant.

## Materials and Methods

### Plant materials and growing conditions

A cassava cultivar (SC8) and friable embryogenic callus (FEC) were cultivated in the dark at 28°C in sterile culture rooms, and *in vivo* cassava plants were inoculated in high-transparency silica glass culture bottles containing Murashige and Skoog basal salt (MS) medium. A photoperiod of 16 h/8 h light/dark condition, with a temperature of 28℃, was adopted. The genetically modified *in vitro* cassava materials were planted in an environment with humidity ranging from 60% to 80%. In the case of tobacco, the optimal culture conditions comprised 14 h of light followed by 10 h of darkness at 24℃.

### Isolation and analysis of the promoter of key enzyme genes in the CG biosynthesis pathway

In this study, the AM560 reference genome of cassava from Phytozome (https://phytozome-next.jgi.doe.gov/) was used, and genomic DNA was extracted from cassava via the cetyltrimethylammonium bromide (CTAB) method. Subsequently, a 2 kb sequence upstream of the *MeCYP79D2* start codon was amplified (Fig. S1). In addition, cis elements within promoter sequences were predicted using the web tool PlantCARE (http://bioinformatics.psb.ugent.be/webtools/plantcare/html/; Table S1).

### Yeast one-hybrid assay

The promoter fragments (−1 to −1505) of *MeCYP79D2* were inserted into the *pAbAi* vector for bait construction. After digestion with *Bst*B I, the *pAbAi-MeCYP79D2* bait vector was used to transform yeast strain Y1HGold and integrated into the yeast genome as reporter strains. The *pAbAi-MeCYP79D2* yeast solution with an optical density (OD)_600_=0.006 was spread evenly on the SD/-Ura medium containing various gradient concentrations of aureobasidin A (AbA, 0–250 ng/mL) to inhibit self-activation of the promoter. A prey vector containing a mixed cDNA library plasmid from cassava storage roots and leaves was introduced to the reporter strains, and growth on the SD/-Ura-Leu (AbA, 300 ng/mL) medium at 30℃ for 3–5 days. The positive colonies were identified via polymerase chain reaction (PCR) with AD detection primer pairs (forward: 5’-CTATTCGATGATGAAGATACCCCACCAAACCC-3’ and reverse: 5’-GTGAACTTGCGGGGTTTTTCAGTATCTACGAT-3’; Table S2).

The point-to-point yeast one-hybrid assay followed the abovementioned steps, and the full-length coding sequence (CDS) of *MePHD1.2* was cloned into the pGADT7 vector. The combination of *pAbAi-P53* and *pGADT7-Rec53* served as a positive control and that of *pAbAi* and *pGADT7-Rec53* as a negative control. The concentration of yeast solution with 0.9% NaCl was diluted to OD_600_=0.6, and a pipette was used to collect 10 µL solution in a 10^−1^ gradient to form a spot on the culture medium. Cultivation was conducted at 30℃ for 3–5 days.

### Phylogenetic tree construction

The AA sequences of *Arabidopsis* and other species proteins, which are similar to those of MePHD1.2, were downloaded from the National Center for Biotechnology Information (https://blast.ncbi.nlm.nih.gov), and a phylogenetic tree was constructed using the neighbour-joining (NJ) method in MAGA6 software.

### Extraction of total RNA and gene expression analyses

Cassava tissue was frozen using liquid nitrogen, and total RNA was extracted using the Plant Total RNA Extraction Kit (Fuji, Chengdu, China). cDNA synthesis was completed using the reverse-transcription reagent MonScript™ RTIII Super Mix with dsDNase (Monad, Suzhou, China). Quantitative real-time PCR (qRT-PCR) was performed using MonAmp™ ChemoHS qPCR Mix (Monad, Suzhou, China) and LightCycler® 480 II (Roche Diagnostics, Germany). Table S2 contains the primer sequences. Each sample underwent three reactions. The relative expression levels of target genes were normalised to that of *Tublin* via the 2^-ΔΔCT^ method.

We collected primary cassava leaf tissues from three individual overexpression (OE) and *cas* transgenic lines (OE-1, OE-2, OE-3, *cas-1*, *cas-2* and *cas-3*), along with a control check (CK) line. Each sample included three independent plants. Then, RNA-seq was performed by GENE DENOVO company (Guangzhou, China). The raw sequence data were deposited to the Genome Sequence Archive (Genomics, Proteomics & Bioinformatics 2021) at the National Genomics Data Center (Nucleic Acids Res 2022), China National Center for Bioinformation, Chinese Academy of Sciences (GSA: CRA014863) (https://ngdc.cncb.ac.cn/gsa).

### Subcellular localisation of MePHD1.2

The *pSL1 Plus* vector containing green fluorescent protein (GFP) was used as the skeleton carrier. The CDS of *MePHD1.2* without a stop codon was fused with the GFP, which is driven by the 35S promoter, to obtain *pSL1 Plus::MePHD1.2-GFP*. Then, the protoplasts were isolated from *in vivo* SC8 cassava leaves of plants grown in an artificial-climate incubator. The cassava leaves were digested using Cellulase R10 (Yakult Honsha, Tokyo, Japan) and Macerozyme R10 (Yakult Honsha, Tokyo, Japan) to prepare the mesophyll cell protoplasts. Polyethylene glycol-mediated cell fusion was used to introduce the nuclear marker mCherry and fusion constructs of *MePHD1.2-GFP* to the cassava mesophyll cell protoplasts. After 12 h incubation in the dark, the protoplasts were harvested, and GFP fluorescence was examined via confocal microscopy.

### Transactivation activity assay

The transcription activation ability was verified using a yeast system, and the full-length CDS of *MePHD1.2* was integrated into the *pGBKT7* vector. Next, the *pGBKT7-MePHD1.2* fusion vector was transformed into Y2HGold yeast strain. We adjusted the OD_600_ of the bacterial liquid to 0.06. The bacterial liquid was subsequently inoculated onto SD/-Trp, SD/-Trp-His, SD/-Trp-Ade and SD/-Trp-His-Ade media. The cultures were incubated at 30℃ for 3–5 days.

Dual luciferase reporter assay was performed to test the transcription activation capability of MePHD1.2. The reconstructed *pGreenII 0800-LUC* vector containing five copies of the GAL4 binding element (5×GAL4) served as the reporter (Wei *et al*., 2017). As the effector, MePHD1.2 was fused with the *GAL4* DNA-binding domain (*GAL4BD*), and an empty *GAL4BD* was used as a control. Tobacco leaves were ground in 1×lysis buffer solution, and the ratio of LUC and REN was determined using the of Dual-Luciferase® Reporter Assay System test kit (E1910, Promega, America). All samples were subjected to more than three repeated measurements.

### Yeast two-hybrid assay

*pGBKT7-MePHD1.2* and *pGADT7* vectors were cotransformed intoY2HGold yeast strain, and the bacterial solution was evenly dispersed on the SD/-Trp-Leu medium. PCR validation involved single colonies and used the AD primer pair. The positive strains were selected for activation. Self-activation validation was carried out on the SD/-Trp-Leu-His-Ade medium with varying concentrations of AbA. Meanwhile, *pGADT7-MePHD1.2* and *pGADT7-MePHD1.1* were cotransformed with the *pGBKT7-MePHD1.2* plasmid into Y2HGold. Protein–protein interactions were determined via colony transfer to the SD/-Trp-Leu-His-Ade medium.

### Dual-Luciferase reporter assay

The promoter fragments (−1 to −1505) of *MeCYP79D2* were inserted into the *pGreenII 0800-LUC* vector as the reporter. The full-length CDS of *MePHD1.2* was cloned into the *pGreenII 62-SK* vector as effector. Along with the *pSoup* helper plasmid, recombinant constructs were introduced to *Agrobacterium* GV3101, followed by their injection into tobacco leaves. Empty *pGreenII 0800* and *pGreenII 62-SK* were used collectively as negative controls. Approximately 48 h later, the tobacco leaves were harvested for measurement of luciferase activity using the same method as the transactivation activity assay.

### Electrophoretic mobility shift assay (EMSA)

The CDS of *MePHD1.2* was cloned into *pMAL-c5X* for in-frame fusion with the maltose binding protein (MBP). The construct of recombinant *MePHD1.2-MBP* was transformed into *Escherichia coli* BL21(DE3). Induction of the corresponding protein expression was completed in 100 mL luria-bertani (LB) liquid culture medium containing 0.6 mmol/L isopropyl β-D-1-thiogalactopyranoside (IPTG). The cells were collected after 12 h induction at 16°C. The process involved the cryogenic ultrasonic lysis of bacteria, followed by protein purification via gel filtration chromatography. Probe synthesis was conducted using the universal primers M13F/R with biotin during PCR, and biotin labelling was applied at the 5′ end by Sangon Biotech company (Shanghai, China). The same but unlabelled DNA fragment was used as a competitor. EMSA was performed using a Light Shift Chemiluminescent EMSA kit (Beyotime, China) and in accordance with the instructions.

### Creation of genetically modified cassava

Gene-specific primers were used to amplify the full-length CDS of *MePHD1.2*, which was subsequently fused with the modified *pCAMBIA1300* plasmid driven by the 35S promoter for OE. The CRISPR/Cas9 system for gene editing was carried by the *pCAMBIA1301* vector. The online website of CRISPR-P v2.0 was used to design target primers that can be accomplished using the (http://crispr.hzau.edu.cn/CRISPR2/). This target can simultaneously target three tandem *MePHDs* in cassava. FEC was applied in a cassava genetic transformation experiment involving mediation by *Agrobacterium* LBA4404. FEC cultivated on Gresshof and Doy Basal Medium (GD) solid medium was inoculated into a fresh one every 20 days. After infection, FEC was transfer to the nylon filter of the GD medium containing carbenicillin (250 mg/L) for 3 weeks. The nylon filter with FECs was transferred to the maturation and germination (MSN) medium containing 20 mg/L hygromycin for 6–8 weeks under a 16 h/8 h photoperiod at 28℃ (the hygromycin content gradually increased from 15 mg/L to 20 mg/L). The medium was refreshed every 2 weeks until the green cotyledons showed maturity. The mature cotyledons were transferred to the shoot-inducting medium cassava embryo maturation (CEM) (which contained 100 mg/L carbenicillin and 10 mg/L hygromycin), which was refreshed every 2 weeks until complete growth of matured sprouts. Afterward, the matured sprouts were cut and transferred to the MS medium.

### Measurement of endogenous CGs

The CGs content was determined using liquid chromatography–tandem mass spectrometry (LC-MS; Schmidt *et al*., 2018).

### Preparation of test solution

Standard stock solution configuration: A total of 10 mg standard substance for linamarin was weighed accurately, dissolved in methanol and diluted to in a 10 mL volumetric flask to prepare a 1 mg/mL standard stock solution of linamarin. A total of 2.5 mg standard substance for lotaustralin was weighed accurately, dissolved in methanol and diluted in a 5 mL volumetric flask to prepare a 0.5 mg/mL lotaustralin-labelled reserve solution.

Standard liquid configuration: Exactly, 100 µL 1 mg/mL linamarin stock solution and 200 µL 0.5 mg/mL lotaustralin stock solution were separately transferred to 10 mL volumetric flasks. These stock solutions were diluted with methanol to obtain a 10 µg/mL mixed standard intermediate solution. A total of 1 mL mixed intermediate liquid was transferred io a 10 mL volumetric flask and diluted with methanol to obtain a mixed solution of 1000 ng/mL, which was used immediately after preparation.

Standard curve configuration: Specifically, 50, 100, 200, 500, 1000, and 5000 µL of the 1000 ng/mL mixed solution were transferred to separate 10 mL volumetric flasks. Each solution was diluted to a certain mark with deionised water and mixed thoroughly. The mixed standard curves of linamarin and lotaustralin were prepared at mass concentrations of 5, 10, 20, 50, 100, 200, and 500 ng/mL.

### Chromatographic conditions

The chromatographic column used was a Shim-pack GIST C18-AQ HP (100 mm×2.1 mm, 1.9 µm). Mobile phase A comprised acetonitrile, and mobile phase B consisted of 0.1% formic acid aqueous solution. Table 1 shows the gradient elution procedure. The flow rate was 0.3 mL/min, the column temperature of the chromatographic column was 0.3℃, and the injection volume was 10 µL.

**Table 1.** Gradient elution program of mobile phase.

| Time (min) | A (%) | B (%) |
| --- | --- | --- |
| 0.5 | 98 | 2 |
| 3 | 93 | 7 |
| 5 | 40 | 60 |
| 6 | 40 | 60 |
| 7 | 98 | 2 |

### MS conditions

An AJS ESI was used as the ion source with positive scanning. Multiple reaction monitoring (MRM) was utilised for CGs detection, with a capillary voltage set at 4000 V, nebuliser pressure at 35 psi and an ion source temperature of 350℃. Nitrogen served as the sheath gas (11.5 L/min) and collision gas (1.65 L/min). Table 2 shows the MRM parameters for the monitoring of each component.

**Table 2.**
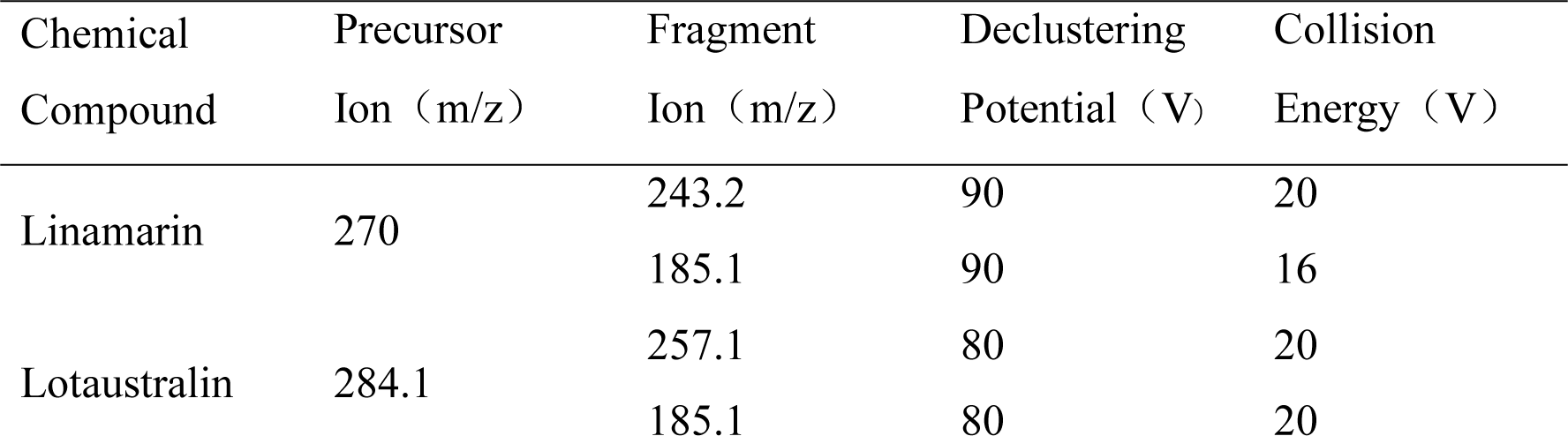
MRM parameter.

### Standard curve establishment

Reference solutions of linamarin and lotaustralin were prepared at varying concentrations. Standard curves were constructed, with the peak area and concentration plotted on the vertical and horizontal axes, respectively. Linear regression equations were derived for linamarin and lotaustralin. The tissue samples, which were ground with liquid nitrogen, were collected from three cassava plants exhibiting gene OE and *cas*, and each measurement was repeated thrice. LC-MS/MS was completed by Hainan Provincial Institute of Food and Drug Inspection. Measurement results were calculated using the following formula: Final concentration×100 mL/Fresh weight (g)×1000.

### Statistical analyses

Each experiment involved at least three independent replicates. The figures represent the mean ± standard error (SE) of at least three independent replicates. Statistical analysis was conducted using the SPSS v.21.0 software, and significance differences among the treatments were assessed using Tukey’s test at P < 0.05 or P < 0.01.

## Results

### Isolation and bioinformatic analysis of MePHD1.2

To explore the regulatory mechanism of CGs biosynthesis in cassava, we cloned and used the 1505 bp promoter of *MeCYP79D2* upstream of its translation initiation site to construct a bait vector. Firstly, the *pAbAi-MeCYP79D2pro* construct was linearised and transformed into competent cells of Y1HGold yeast, and growth vigour of the recombinant strain with *pAbAi-MeCYP79D2pro* was effectively inhibited on the SD/-Ura medium containing 250 ng/ml AbA (Fig. 2a). Secondly, a mixed cDNA library of storage roots, stems and leaves was used as prey to perform the Y1H assay, and 1103 positive clones were screened out. A total of 109 protein-coding genes, including 32 transcription factors, were identified after sequencing and functional annotation (Table S3 and Fig. 2b). Among the candidates, the PHD member (Manes.12G071000) appeared most frequently and accounted for approximately 5% of the total positive clones.

**Fig. 2.**
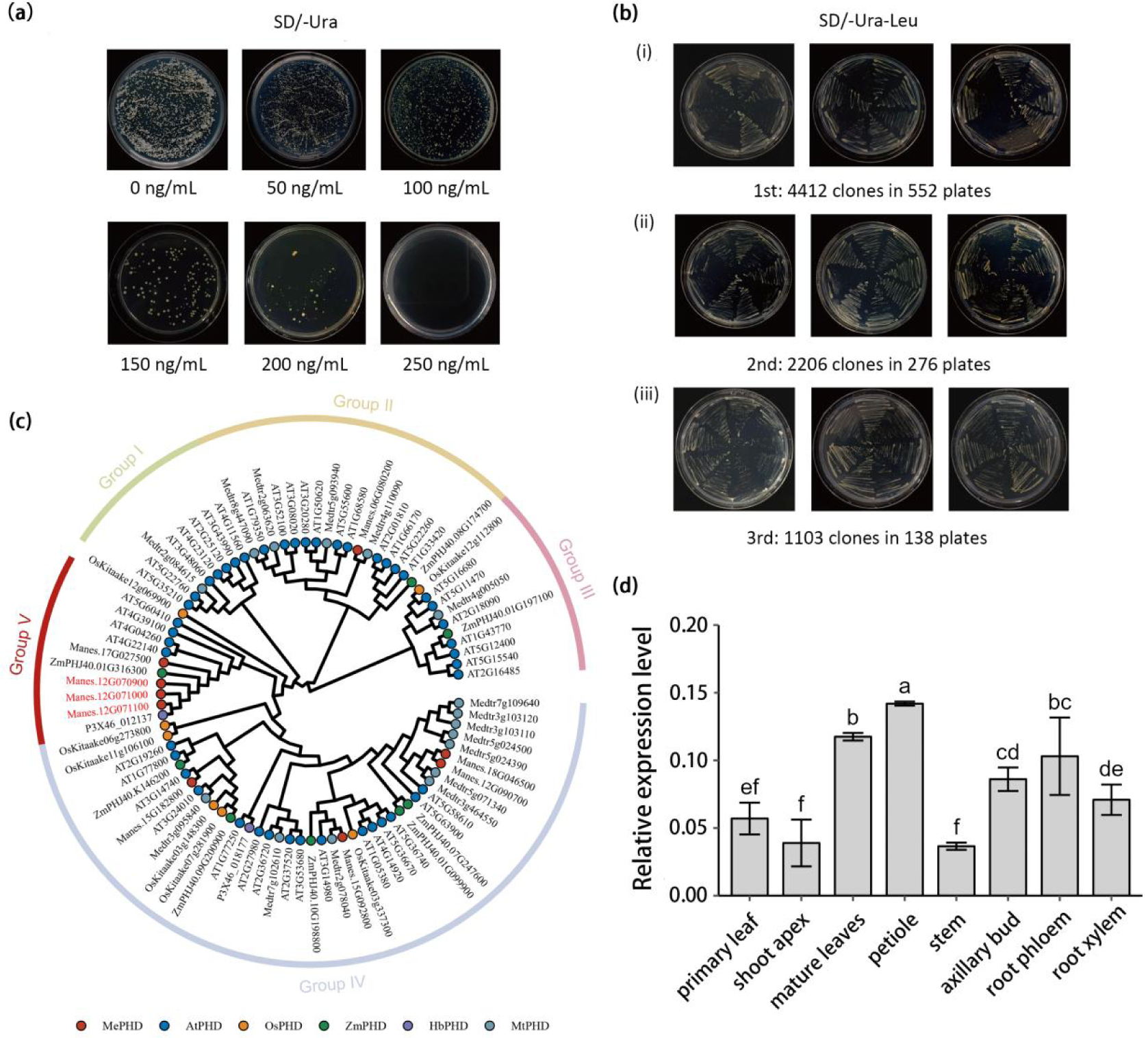
Yeast one-hybrid assay and characterisation of PHD transcription factors. (a) Inhibition of the self-activation of *pAbAi-MeCYP79D2*. AbA was added to SD/-Ura medium at concentrations of 50, 100, 150, 200 and 250 ng/mL. (b) Verify 3 times to confirm the positive clones. (c) Phylogenetic analysis of homologous PHD transcription factors from various species. AA sequences were aligned using MEGA6 software, and a phylogenetic tree was constructed via the NJ method. MePHD/cassava, AtPHD/arabidopsis, OsPHD/rice, ZmPHD/maize, HbPHD/rubber tree and MtPHD/alfalfa. (d) Relative expression levels of *MePHD1.2* in various tissues of SC8 cassava variety. The data are the means ± standard deviation (SD) (n=3). Statistically significant differences were determined via one-way analysis of variance (ANOVA). Different letters indicate statistically significant differences.

The cassava genome contains 11 members of PHD family (Fig. S2a), with two of the members arranged sequentially with Manes.12G071000 on chromosome 12. The AA sequences of two other members exhibit high homology and differ only in terms of specific AA residues at the C-terminal (Fig. S2b). These members were named MePHD1.1 (Manes.12G070900), MePHD1.2 (Manes.12G071000) and MePHD1.3 (Manes.12G071100) based on their order on Chromosome 12.

PHD member is a zinc finger protein widely present in eukaryotes and belongs to the zinc finger domain family; this protein performs a crucial function in transcription and chromatin state regulation. MePHD1.2 presents a relatively high homology with chromatin remodelling proteins, such as OsPHD, ZmPHD, HbPHD, MtPHD, etc. and the highest homology with AtEBS (At4G22140) in *Arabidopsis* (Fig. 1c). *MePHD1.2* encodes 216 AA residues, and its protein comprises one BAH domain and one PHD domain with two Zn-binding sites (Fig. S2c). *MePHD1.2* is also expressed in all tissues of cassava, with relatively high expression levels observed in mature leaves, axillary buds and roots (Fig. 2d).

### Characterisation of transcription factor MePHD1.2

*pCAMBIA1300-MePHD1.2-GFP* was transformed into cassava protoplasts for transient expression, and an intense fluorescence of the GFP, which indicates localisation of MePHD1.2 in the nucleus, was observed in the cell nucleus through confocal laser microscopy (Fig. 3a). In addition, MePHD1.2 was located in the nucleus of tobacco leaves (Fig. S3a). Thus, MePHD1.2 is a nuclear localised protein.

**Fig. 3.**
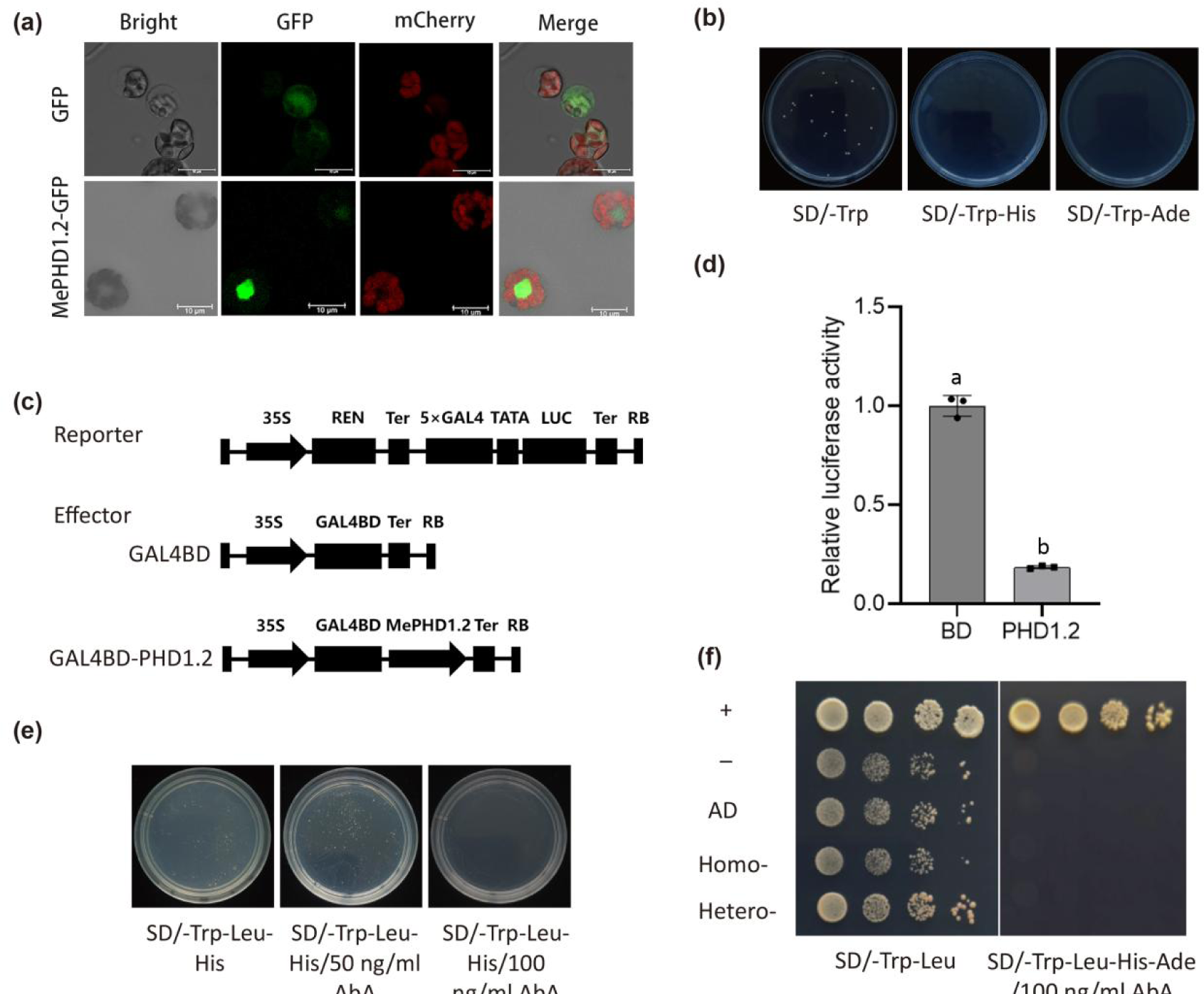
MePHD1.2 is a transcription factor with a transcriptional inhibitory activity. (a) Subcellular localisation of MePHD1.2. Full-length MePHD1.2 fused with the improved GFP and the nucleus marker mCherry were coexpressed in cassava protoplast cells. *pSL1 Plus-GFP* served as a negative control. Scale bar: 10 µm. (b) Validation of MePHD1.2 transcription activation activity in yeast cells. The *pGBKT7-MePHD1.2* recombinant plasmid was transformed into Y2HGold competent cells, and the bacterial solution was diluted to OD_600_=0.002. A 100 µL yeast liquid was obtained and subsequently applied to the SD/-Trp, SD/-Trp-His and SD/-Trp-Ade media. The plates were incubated at 30℃ for 2–3 days in an incubator. (c) Transcription activation activity reporting system model. (d) Relative luciferase activity was detected in the cassava protoplast. The data are the mean ± SD (n=3). One-way ANOVA revealed statistically significant differences. Different letters denote statistically significant differences. (e) Inhibition of the self-activation of MePHD1.2 transcription factor. (f) Oligomerisation analysis of MePHD1.2 transcription factor. Cotransformation of *pGBKT7-MePHD1.2* with homologous *pGAD-MePHD1.2* and heterologous *pGAD-MePHD1.1* was conducted in yeast Y2HGold, followed by culture on SD/-Trp-Leu-His-Ade media containing 100 ng/mL AbA. ‘+’ represents the combination of *pGBKT7-53* and *pGADT7*, ‘-’ denotes the combination of *pGBKT7-Lam* and *pGADT7*, ‘AD’ indicates the combination of *pGBKT7-MePHD1.2* and *pGADT7*, ‘Homo –’ refers to homologous interaction and ‘Hetero –’ means heterologous interaction.

To determine whether MePHD1.2 possesses transcriptional activation capability, we coated *pGBKT7-MePHD1.2* yeast strains on SD/-Trp, SD/-Trp-His and SD/-His-Ade media. However, only the positive recombinant yeast strain with *pGBKT7-53* + *pGADm* grew well on the SD/-Trp-His and SD/-His-Ade medium (Fig. 3b). Meanwhile, the yeast strains with six truncated *MePHD1.2* segments on *pGBKT7* failed to grow on SD/-Trp-His-Ade (Figs. S3b–3c). The finding proves that MePHD1.2 lacks a transcriptional activation activity.

The *LUC* reporter gene was fused with the *5×GAL4* DNA-binding element + TATA box, with *35S::REN* as the internal reference and *GAL4BD* and *GAL4BD-PHD1.2* as effectors to perform a dual-luciferase reporter assay (Fig. 3c). *GAL4BD* exhibited a LUC/REN value 5.4 times that of *GAL4BD-PHD1.2*, which indicates that MePHD1.2 considerably reduced the expression of *LUC* reporter gene (Fig. 3d). This finding proves the transcriptional inhibitory activity of MePHD1.2.

Numerous transcription factors perform their functions through the formation of homodimers or heterodimers. The *pGBKT7-MePHD1.2* and *pGADm* constructs were cotransformed into Y2HGold yeast competent cells. Incremental concentrations of AbA were added to the yeast culture medium, and self-activation of MePHD1.2 was inhibited when the concentration of AbA reached 100 ng/mL (Fig. 3e). Next, oligomerisation analysis of MePHD1.2 was conducted, and the results show that MePHD1.2 cannot form homodimers nor heterodimers, which indicates its independent mode of function (Fig. 3f).

### MePHD1.2 directly binds to the PD2 segment and represses the transcriptional activity of *MeCYP79D2*

In the examination of whether MePHD1.2 binds to the *MeCYP79D2* promoter, the yeast cells cotransformed with *pAbAi-MeCYP79D2pro* and *pGADT7-MePHD1.2* grew in the SD/-Ura-Leu medium supplemented with 300 ng/mL AbA (Fig. 4a), which suggests that MePHD1.2 can recognise and bind to the *MeCYP79D2* promoter. The 1505 bp promoter of *MeCYP79D2* was truncated into three segments, with each measuring approximately 500 bp, and named PD1, PD2 and PD3 (Fig. 4b). PD1/2/3 were used to construct bait vectors for the Y1H assay with *MePHD1.2* as prey. Firstly, only the yeast strain that contain *pAbAi-PD2* and *pGADT7-MePHD1.2* can grow on the medium SD/-Ura-Leu supplemented with 50 ng/mL AbA, which implies the possible binding of MePHD1.2 to the PD2 segment of the *MeCYP79D2* promoter (Fig. 4c).

**Fig. 4.**
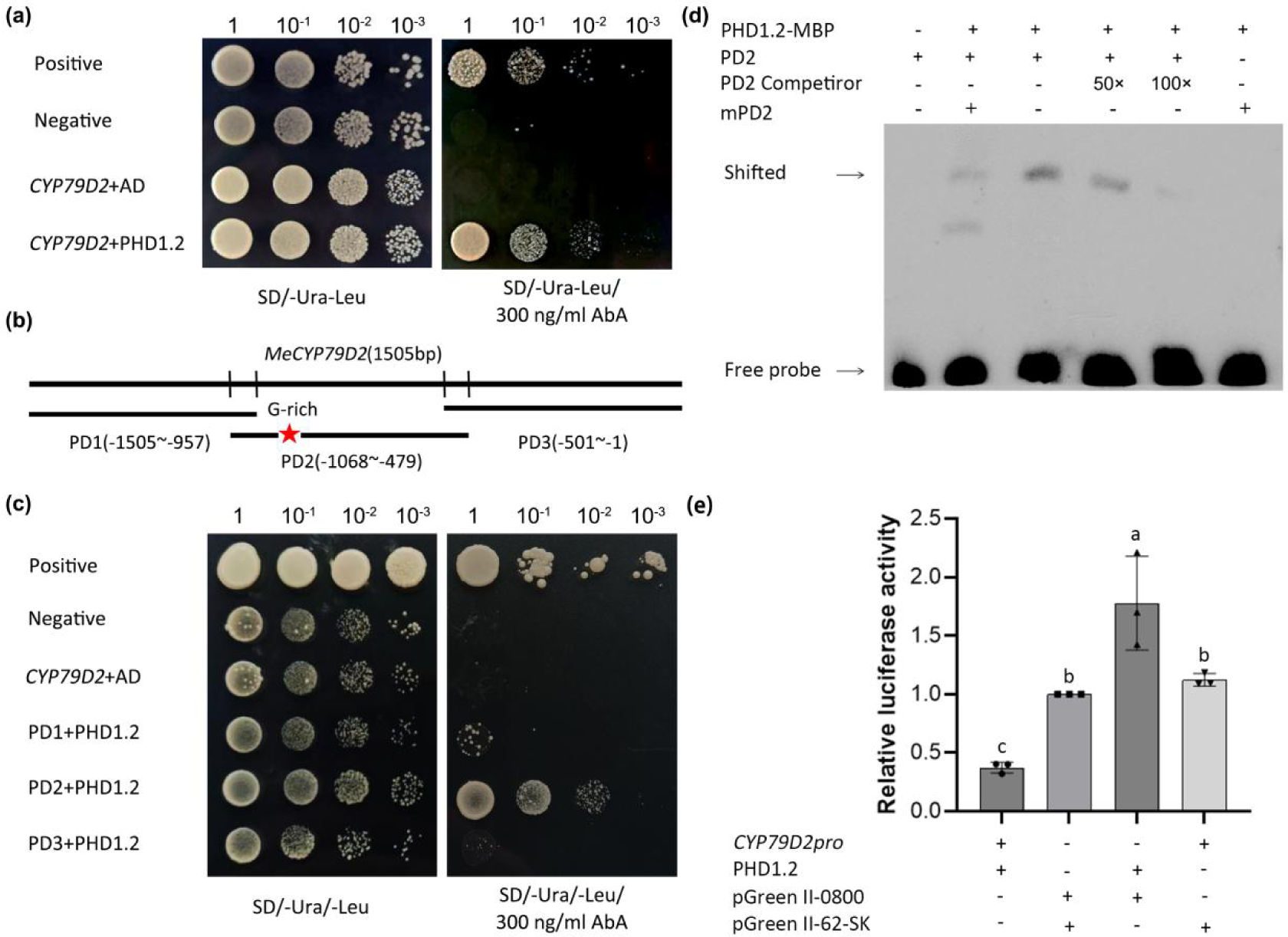
Binding of MePHD1.2 to the PD2 segment of *MeCYP79D2* promoter and its repressive action on transcription activity. (a) Yeast one-hybrid assay between MePHD1.2 and *MeCYP79D2* promoter*. pGADT7-MePHD1.2* was transformed into yeast competent cells containing *pAbAi-MeCYP79D2pro*. The cells were cultured at various gradients of bacterial concentrations on SD/-Ura-Leu medium supplemented with 300 ng/mL AbA. Positive and negative controls represented the combinations of *pAbAi-p53* and *pGADT7* and that of *pAbAi* and *pGADT7*, respectively. AD indicates *pGADT7*. (b) Pattern diagram of truncated promoter segments. PD1, PD2 and PD3 represent the three segments of *MeCYP79D2*. (c) Yeast one-hybrid assay between MePHD1.2 and PD1/2/3. (d) EMSA of MePHD1.2 bound to the PD2 segment *in vitro*. Biotin-labelled DNA probes (PD2 or mPD2) were incubated with the MBP-MePHD1.2 protein, and DNA–protein complexes were separated on 6% native polyacrylamide gels. The cold probes of the competitor were the unlabelled probe; ‘50×’ and ‘100×’ represent 50- and 100-fold molar excess, respectively. The symbol ‘+’ or ‘-’ indicates the presence or absence of the corresponding proteins and probes, respectively. (e) Dual-luciferase assay. The LUC/REN ratio, which was set to 1, was normalised to the samples combined with *pGreenII 62-SK* and *pGreenII 0800-LUC*. LUC, firefly luciferase activity; REN, Renilla luciferase activity. The data are the mean ± SD (n=3). Statistically significant differences were determined using one-way ANOVA. Different letters denote statistically significant differences.

Secondly, PD2 was labelled with 5′-biotin for the EMSA. The shift in band was detected when PD2-biotin was incubated with MePHD1.2-MBP fusion protein. However, no band shift occurred with MBP protein alone. The density of band shifts gradually decreased with the increase in the concentration of the unlabelled competitor PD2 (Fig. 4d).

Thirdly, to further clarify the regulatory effect of MePHD1.2 on the transcription of *MeCYP79D2*, we used dual-luciferase assay to determine the relative luciferase activity in tobacco leaves. The three negative controls had LUC/REN values 2.7 to 4.7 times higher than those of the cotransformation with MePHD1.2 effector and *MeCYP79D2pro* reporter (Fig. 4e). Thus, MePHD1.2 can bind to the PD2 segment of *MeCYP79D2* promoter and negatively regulate its transcription.

### MePHD1.2 represses the expression of *MeCYP79D2* while increases the CGs content in *mephd1.2* mutant leaves

To assess the physiological function of MePHD1.2, we developed 40 OE lines and verified them by rooting test (Fig. S4a). Lines OE1, OE2 and OE3, which had one-copy T-DNA according to Southern blotting and genome re-sequencing, were selected (Figs. 5a–5b). These three OE lines had relatively high expressions of *MePHD1.2* compared with the other OE lines, as shown by qRT-PCR assay (Fig. S4b). Moreover, Hi-TOM assay identified 36 gene-edited lines as homozygous mutant, and three (*cas1*, *cas2* and *cas3*) were selected because of the presence of their monoallelic homozygous mutant at the target site (Figs. 5c and S4c).

**Fig. 5.**
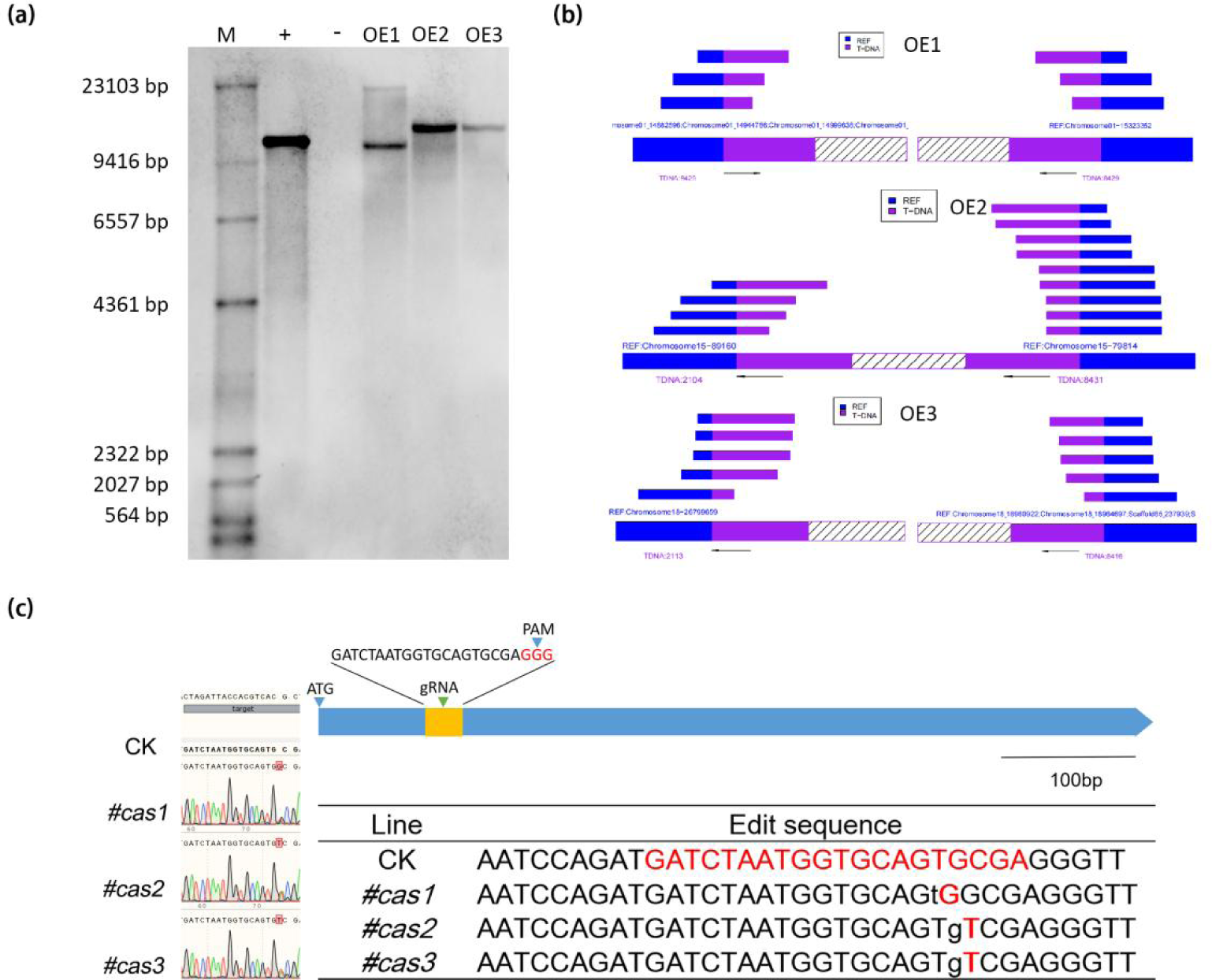
Identification of OE and *cas* lines. (a) Southern blotting of *MePHD1.2* OE lines. (b) Genome resequencing alignment of *MePHD1.2* OE lines. (c) Mutant types of *MePHD1.2 cas* lines. CK represents the regenerated plant of SC8 infected with an empty vector in *Agrobacterium*.

To clarify the regulatory effect of MePHD1.2 on CG biosynthesis pathway, we determined the contents of linamarin and lotaustralin in the first unfolded leaves of OE1/2/3, *cas1/2/3* lines and CK via LC-MS. MS revealed two distinct peaks. The first compound was linamarin, followed by lotaustralin (Fig. S5). The standard curve equation of linamarin is y=28.262x+2143.796, R^2^=0.997; that of lotaustralin is y=40.020x+1569.524, R^2^=0.989 (Fig. S6). The average contents of linamarin and lotaustralin in the *cas* lines reached 454.02±18.60 and 85.20±23.85 mg/kg, respectively, which are significantly higher than those in CK (Figs. 6a–6b). However, the contents of these CGs showed no significant difference between OE lines and CK. In addition, the expression levels of *MeCYP79D2* in the *cas* lines were significantly higher than that in CK, but the differences between the OE lines and CK were not significant (Fig. 6c). Significant differences were observed in the gene expression levels between OE and *cas* lines in terms of the CG metabolism pathway. The CG biosynthesis genes (*MeCYP79D2*, *MeCYP71E7* and *MeUDT85K4/5*), CG decomposition genes (*MeBGLU12A/B* and *MeHNL)* and CGs reutilisation genes (*MeCAS1a/b* and *MeNIT4A/B)* were all up-regulated in the *cas* lines (Figs. 6d–6f). Thus, the functional deficiency of MePHD1.2 in *cas* lines weakened its transcription inhibition of *MeCYP79D2*, which led to an up-regulation of various CG genes and thus improved the biosynthesis efficiency of CGs in the leaves.

**Fig. 6.**
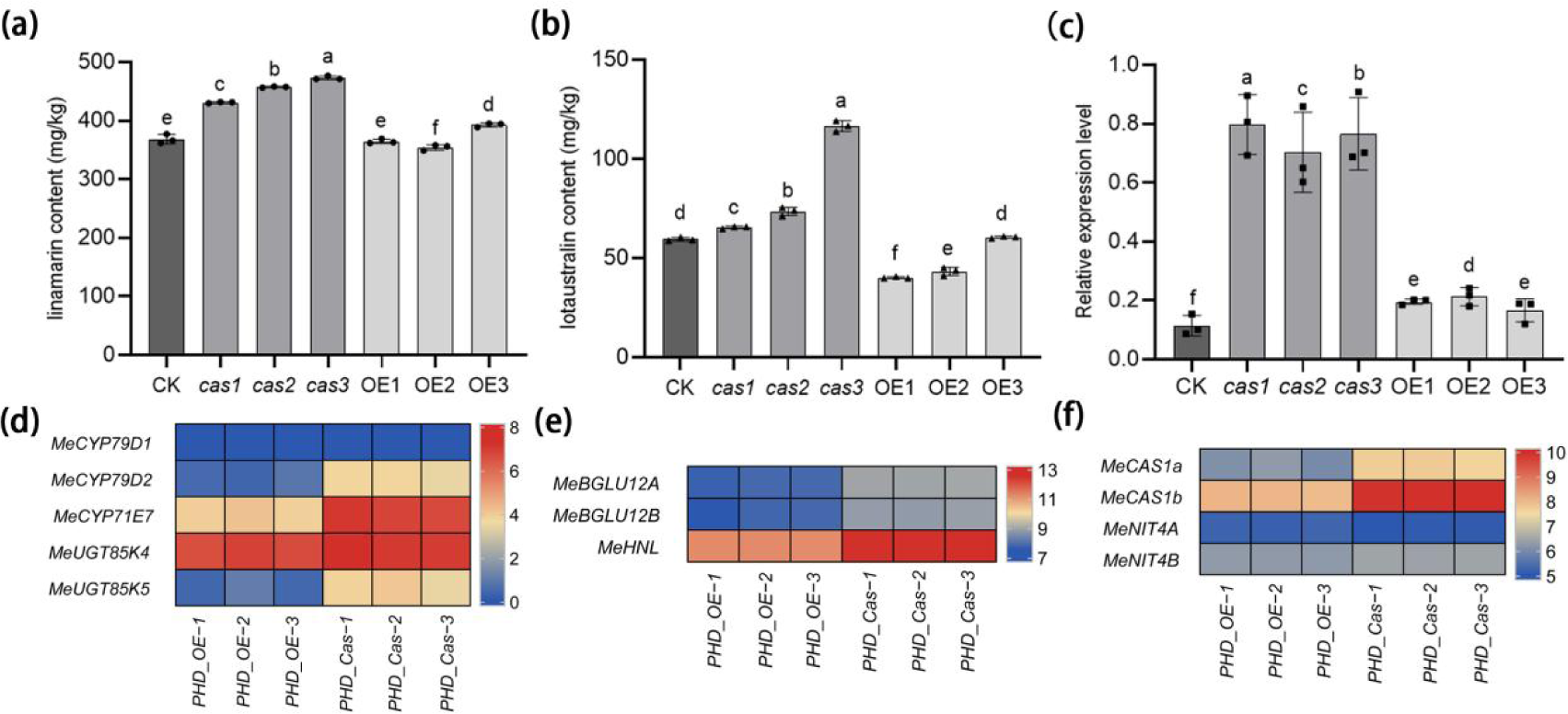
Determination of CG content and relative expression levels of *MeCYP79D2*. (a–b) Linamarin and lotaustralin contents in CK and *cas*/OE lines. (c) Expression levels of *MeCYP79D2* in CK and *cas*/OE lines. (d–f) Expression differences of key genes related to CG biosynthesis, decomposition and reutilisation between OE and *cas* lines. The data are the mean ± SD (n=3). One-way ANOVA revealed statistically significant differences. Different letters indicate statistically significant differences.

## Discussion

### MePHD1.2 can directly bind to the *MeCYP79D2* promoter and negatively regulate its transcriptional activity

PHD is a homologous domain protein with zinc fingers, and it can regulate the transcriptional activity of its targets by binding to the G-rich (GNGGTG) element in their promoters. Soybean GmPHDs bind to the GTGGAG element and improve stress resistance capability (Winicov *et al*., 1999; Wei *et al*., 2009 and 2015). Banana MaPHD1 can also directly bind to the G-rich element in *MaXTH6* promoter, which is a gene related to cell-wall degradation, and down-regulate its transcriptional activity to delay fruit ripening(Wei *et al*., 2017). Consistently, in this study, MePHD1.2 can directly bind to the PD2 segment of *MeCYP79D2* promoter, a G-rich element located in the PD2 segment, and negatively regulate its transcriptional activity (Figs. 4d–4e).

PHD proteins also act as readers to recognise histone modifications; they can bind to the ‘tails’ of histone H3 with various modifications to recognise the histone code, which results in changes in histone recognition preference to inhibit or activate DNA transcription (Narro-Diego *et al*., 2017; Miura *et al*., 2020). At low temperatures, vernalisation induces a state transition in flowering-related genes, such as the floral integrators *FLOWERING LOCUS T* (*FT*) and *FLOWERING LOCUS C* (*FLC*), via PHD-mediated chromatin remodelling, which involves histone dimerisation and trimethylation. Alterations in histone methylation levels can either promote or inhibit the flowering of *Arabidopsis* (Greb *et al*., 2007; Lopez-Gonzalez *et al*., 2014). The PHD-type protein AL transforms the chromatin state of seed development genes, namely, *ABSCISIC ACID INSENSITIVE 3* (*ABI3*) and *DELAY OF GENATION 1* (*DOG1*), through the formation of a PHD-polycomb reactive complex 1 (PRC1) complex with PRC1. Compression of chromatin causes inhibition in the transcription activities of *ABI3* and *DOG1*, which promotes seed germination (Molitor *et al*., 2014). The RNA silencing effect of the gene *ARGONAUTE 5* (*AGO5*) is inhibited by the BAH-PHD-CPL2 complex (BPC), which results from the binding of BAH-PHD to CPL2, a plant-specific Pol II carboxyl terminal domain phosphatase. This complex enables CPL2-mediated CTD dephosphorylation to inhibit the release of Pol II from the transcription start site, which results in a suppressed gene transcription (Zhang *et al*., 2020). Further research is required to ascertain whether this complex can recognise histones or recruit coregulatory factors to cooperatively regulate the transcription of its targets.

### MePHD1.2 regulates CG *synthesis* by inhibiting the transcriptional activity of *MeCYP79D2*

The variation in key AA sites of the transcription factor basic helix-loop-helix (bHLH) in *Lotus Japonicus* and almond prevents them from forming dimers or affects their direct recognition of promoters of downstream genes involved in CG synthesis. This condition leads to a weakened or lost function, reduced transcriptional activities of genes encoding the rate-limiting enzymes involved in CG biosynthesis and a consequent decrease in the CG content (Sánchez-Pérez, 2019; Chen *et al*., 2022). Studies on cassava have shown that the down-regulation of *MebHLH72* and *MebHLH114* can lead to a decrease in the linamarin content of cassava leaves (An *et al*., 2022). In addition, OE of bHLH transcription factors *TSAR1* or *TSAR2* in *Alfalfa,* causes the increase in the expression levels of genes involved in the biosynthesis of glycosides /triterpenoid saponin and rapidly increases the efficiency of their synthesis (Mertens, *et al*., 2016). In this study, significantly increased expression of *MeCYP79D2* and CG contents in leaves were observed in the *cas* lines of MePHD1.2 compared with those in CK. However, the difference between the OE lines and CK was not significant. Based on these findings, knockout of *MePHD1.2* probably reduced the degree of methylation in the *MeCYP79D2* promoter and prevented the recruited coregulators from binding to the promoter, which inhibited their capability to suppress transcriptional activity.

### CG accumulation may grant cassava the ability to tolerate barren conditions

Cassava exhibits resilience to barren conditions and can achieve a certain storage root yield in soils where other crops experience failure in completing their life cycle (Nguyen *et al*., 2001). Plants biosynthesise various secondary metabolites through primary metabolites to perform various physiological functions; when plants suffer from nutritional stress or are in a critical developmental cycle, these secondary metabolites can be decomposed and assimilated back as primary metabolites to deal with the new condition (Sugiyama *et al*., 2021). CGs are nitrogen-containing secondary metabolites that can be decomposed into cyanohydrins and HCN under the continuous action of linamarase and hydroxynitrile hydrolase (HNL). HCN and cysteine are used to synthesise cyanalanine, which is a compound catalysed by cyanalanine synthase (CAS). Finally, cyanalanine is decomposed into Asp and NH_3_ through catalysis by nitrile hydrolase (NIT4) (Picmanova *et al*., 2015; Nielsen *et al*., 2016). CGs play an important role in the resistance to herbivorous and insect infestations and act as a form of nitrogen reserve for the increased content of reduced nitrogen (Gladow and Woodrow, 2000). RNAi of *CYP79D1/D2* gene considerably reduces cyanoside content in cassava; RNAi plants cannot complete their life cycle in an environment that lacks reduced nitrogen (Siritunga and Sayre, 2003). The OE of *HNL* increased the contents of Asp and other AAs in cassava roots (Siritunga *et al*., 2004; Leyva-Guerrero *et al*., 2012). Therefore, CG decomposition can increase the supply of AAs and reduced nitrogen (Narayanan *et al*., 2011; Zidenga *et al*., 2017). CGs are mainly synthesised in the leaves and transported to storage roots in cassava. Nitrogen deficiency stimulates the formation and development of lateral roots to increase nutrient supply (Zhang and Forde, 2000). It was speculated that the transport of CGs from leaves to the roots will further strengthened, and decompose into AAs and the reduced nitrogen to supply root development under low nitrogen stress. Altogether, the increase in CG content of the leaves of *mephd1.2* mutant maybe improve its ability to tolerate nitrogen deficiency.

### New cassava varieties with high CG content in leaves and low CG in storage roots can be bred with the assistance of gene editing and molecular markers

Phloem girdling test revealed the accumulation of CGs in the upper part of the girdling incision and their decreased levels in the lower part, which indicates that CGs in cassava roots are primarily transported from the leaves (Kannangara *et al*., 2011; Gleadow *et al*., 2014; Jørgensen *et al*., 2023). A weakened transport capacity can considerably reduce the content of CGs in cassava storage roots (Lieberman *et al*., 2024). Genome-wide association analysis identified two major loci influencing CG content in cassava storage roots. The multidrug and toxic compound extrusion (*MATE*) gene on chromosome 16 contributed 30% of the CG content variation in experimental populations. The G→A mutation in the fourth exon of *MATE* gene changes the original encoded Ala to Thr, which disrupts the transmembrane structure of MATE protein and substantially reduces the transport efficiency of CGs (Schmidt *et al*., 2018; Ogbonna *et al*., 2021). The high CG content in leaves of *MePHD1.2* mutant and inefficient CG transport characteristics of parent lines can be utilised in cross-breeding and provide assistance in the molecular identification of genotypes for the rapid development of new cassava varieties with high CG content in leaves and low CGs in storage roots. Furthermore, the inclusion of these cassava varieties in agricultural production will not only improve the pest resistance of the plant and reduce the application of chemical pesticides, but also enable the production of edible cassava with low CG content in storage roots. Such achievement will be helpful in the green production of cassava and ensure food safety for populations in Asia, Africa, and Latin America.

## Acknowledgements

This work was supported by grants from the China Agriculture Research System (CARS-11) and the Chinese Academy of Tropical Agricultural Sciences for the Science and Technology Innovation Team of the National Tropical Agricultural Science Center (CATASCXTD202301). Additional support was provided by the Hainan Province Graduate Innovation Research Project (Hyb2020-09) and the Key Laboratory of Tropical Fruits and Vegetables Quality and Safety for State Market Regulation (KF-2023016). We express our gratitude to Wei Wei and Shengyan Li for providing the *LUC* reporter gene with the *5×GAL4* DNA-binding element + TATA box and the *GAL4BD* vector.

## Author contributions

ML, YL, and XC conceptualized and designed the experiments. ML created the CRISPR-Cas9 and overexpression lines, while ML and YJL collaborated on the manuscript design and writing. ML and XZ conducted the experiments, with XYZ, QL and QY focusing on CGs content analysis. WM assisted with data analysis. WW, JL and XC originated the project idea and provided supervision throughout. All authors have reviewed and approved the final version of the manuscript.

## Data availability

All data supporting the findings are included in this manuscript. Additional supporting informations can be found in the Supporting Information section at the end of the article.

## Supporting Information

Additional supporting information may be found online in the Supporting Information section at the end of the article.

**Fig. S1** Promoter sequence (2 kb) of *MeCYP79D2* gene.

**Fig. S2** Characterisation of MePHD1.2 in cassava.

**Fig. S3** Subcellular localisation and transcription activation region assay of MePHD1.2.

**Fig. S4** Identification and phenotypic characterisation of MePHD1.2 transgenic plants.

**Fig. S5** MS peak of linamarin and lotaustralin determination.

**Fig. S6** Standard curves of linamarin and lotaustralin in LC-MS/MS detection.

**Table S1** Part of cis element on the MeCYP79D2 promoter.

**Table S2** Primers used for vector construction in this study. Table S3 Partially interacting proteins identified via Y1H assay.

## References

An F, X Xh, Chen T, Xue JJ, Luo XQ, Ou WJ, Li KM, Cai J, Chen SB. 2022. Systematic analysis of bHLH transcription factors in cassava uncovers their roles in postharvest physiological deterioration and cyanogenic glycosides biosynthesis. Frontiers in Plant Science 13: 901128.

Andersen MD, Busk PK, Svendsen I, Møller BL. 2000. Cytochromes P-450 from cassava (*Manihot esculenta* Crantz) catalyzing the first steps in the biosynthesis of the cyanogenic glucosides linamarin and lotaustralin: cloning, functional expression in pichia pastoris, and substrate specificity of the isolated recombinant enzymes. Journal of Biological Chemistry 275:1966–1975.

Bak S, Kahn RA, Nielsen HL, Møller BL, Halkier BA. 1998. Cloning of three A-type cytochromes P450, CYP71E1, CYP98, and CYP99 from *Sorghum bicolor* (L.) Moench by a PCR approach and identification by expression in *Escherichia coli* of CYP71E1 as a multifunctional cytochrome P450 in the biosynthesis of the cyanogenic glucoside dhurrin. Plant molecular biology 36: 393–405.

Ballhorn DJ, Kautz S, Laumann JM. 2016. Herbivore damage induces a transgenerational increase of cyanogenesis in wild lima bean (*Phaseolus lunatus*). Chemoecology 26:1–5.

Banea-Mayambu JP, Tylleskär T, Gitebo N, Matadi N, Gebre-Medhin M, Rosling H. 1997. Geographical and seasonal association between linamarin and cyanide exposure from cassava and the upper motor neurone disease konzo in former Zaire. Tropical Medicine & International Health 2:1143–1151.

Chen C, Liu F, Zhang K, Niu X, Zhao H, Liu Q, Georgiev MI, Xu X, Zhang X, Zhou M. 2022. MeJA-responsive bHLH transcription factor LjbHLH7 regulates cyanogenic glucoside biosynthesis in *Lotus japonicus*. Journal of experimental botany 18: 2650–2665.

Dudareva N, Pichersky E, Gershenzon J. 2004. Biochemistry of plant volatiles. Plant physiology 135:1893–902..

Easson ML, Malka O, Paetz C, Hojná A, Reichelt M, Stein B, van Brunschot S, Feldmesser E, Campbell L, Colvin J, Winter S. 2021. Activation and detoxification of cassava cyanogenic glucosides by the whitefly *Bemisia tabaci*. Scientific reports 24: 13244.

Fischbach MA, Clardy J. 2007. One pathway, many products. Nature chemical biology 3:353–355.

Forslund K, Morant M, Jørgensen B, Olsen CE, Asamizu E, Sato S, Tabata S, Bak S. 2004. Biosynthesis of the nitrile glucosides rhodiocyanoside A and D and the cyanogenic glucosides lotaustralin and linamarin in *Lotus japonicus*. Plant physiology 135: 71–84.

Ganjewala D. 2010. Advances in cyanogenic glycosides biosynthesis and analyses in plants: A review. Acta Biologica Szegediensis 54: 1–4.

Gleadow RM, Møller BL. 2014. Cyanogenic glycosides: synthesis, physiology, and phenotypic plasticity. Annual review of plant biology 65: 155–85.

Gleadow RM, Woodrow IE. 2000. Temporal and spatial variation in cyanogenic glycosides in Eucalyptus cladocalyx. Tree Physiology 20: 591–598.

Gomez MA, Berkoff KC, Gill BK, Iavarone AT, Lieberman SE, Ma JM, Schultink A, Karavolias NG, Chauhan RD, Taylor NJ, Staskawicz BJ. 2023. CRISPR-Cas9-mediated knockout of *CYP79D1* and *CYP79D2* in cassava attenuates toxic cyanogen production. Frontiers in Plant Science 13: 1079254.

Greb T, Mylne JS, Crevillen P, Geraldo N, An H, Gendall AR, Dean C. 2007. The PHD finger protein VRN5 functions in the epigenetic silencing of Arabidopsis FLC. Current biology 17: 73–78.

Jain K, Fraser CS, Marunde MR, Parker MM, Sagum C, Burg JM, Hall N, Popova IK, Rodriguez KL, Vaidya A, Krajewski K. 2020. Characterization of the plant homeodomain (PHD) reader family for their histone tail interactions. Epigenetics & chromatin 13: 1–11.

Jørgensen K, Bak S, Busk PK, Sørensen C, Olsen CE, Puonti-Kaerlas J, Møller BL. 2005. Cassava plants with a depleted cyanogenic glucoside content in leaves and tubers. Distribution of cyanogenic glucosides, their site of synthesis and transport, and blockage of the biosynthesis by RNA interference technology. Plant Physiology 139: 363–374.

Jørgensen K, Morant AV, Morant M, Jensen NB, Olsen CE, Kannangara R, Motawia MS, Møller BL, Bak S. 2011. Biosynthesis of the cyanogenic glucosides linamarin and lotaustralin in cassava: isolation, biochemical characterization, and expression pattern of CYP71E7, the oxime-metabolizing cytochrome P450 enzyme. Plant physiology 155: 282–292.

Kannangara R, Motawia MS, Hansen NK, Paquette SM, Olsen CE, Møller BL, Jørgensen K. 2011. Characterization and expression profile of two UDP-glucosyltransferases, UGT85K4 and UGT85K5, catalyzing the last step in cyanogenic glucoside biosynthesis in cassava. The Plant Journal 68: 287–301.

Kongsawadworakul P, Viboonjun U, Romruensukharom P, Chantuma P, Ruderman S, Chrestin H. 2009. The leaf, inner bark and latex cyanide potential of Hevea brasiliensis: evidence for involvement of cyanogenic glucosides in rubber yield. Phytochemistry 70: 730–739.

Lai D, Maimann AB, Macea E, Ocampo CH, Cardona G, Pičmanová M, Darbani B, Olsen CE, Debouck D, Raatz B, Møller BL. 2020. Biosynthesis of cyanogenic glucosides in Phaseolus lunatus and the evolution of oxime-based defenses. Plant Direct 4: e00244.

Latif S, Müller J. 2015. Potential of cassava leaves in human nutrition: A review. Trends in Food Science & Technology 44: 147–158.

Lebot V. 2019. Tropical root and tuber crops. Cabi.

Leyva-Guerrero E, Narayanan NN, Ihemere U, Sayre RT. 2012. Iron and protein biofortification of cassava: lessons learned. Current opinion in biotechnology 23: 257–264.

Lieberman SE, Gueorguieva GA, Gill BK, Litvak L, Cruz AG, Lyons JB, Cho MJ, Karavolias N. 2024. Transporter editing in cassava indicates local production of cyanogenic glucosides in, and export from, cassava roots. Plant Biotechnology Journal 22: 790.

López-González L, Mouriz A, Narro-Diego L, Bustos R, Martínez-Zapater JM, Jarillo JA, Piñeiro M. 2014. Chromatin-dependent repression of the Arabidopsis floral integrator genes involves plant specific PHD-containing proteins. The Plant Cell 26: 3922–3938.

Ma PA, Chen X, Liu C, Xia ZQ, Song Y, Zeng CY, Li YZ, Wang WQ. 2018. MePHD1 as a PHD-finger protein negatively regulates ADP-glucose pyrophosphorylase small subunit1a gene in cassava. International Journal of Molecular Sciences 19: 2831.

McMahon J, Sayre R, Zidenga T. 2022. Cyanogenesis in cassava and its molecular manipulation for crop improvement. Journal of Experimental Botany 73: 1853–1867.

Mertens J, Pollier J, Vanden Bossche R, Lopez-Vidriero I, Franco-Zorrilla JM, Goossens A. 2016. The bHLH transcription factors TSAR1 and TSAR2 regulate triterpene saponin biosynthesis in *Medicago truncatula*. Plant physiology 170: 194–210.

Miura K, Renhu N, Suzaki T. 2020. The PHD finger of *Arabidopsis* SIZ1 recognizes trimethylated histone H3K4 mediating SIZ1 function and abiotic stress response. Communications biology 3: 23.

Molitor AM, Bu Z, Yu Y, Shen WH. 2014. *Arabidopsis* AL PHD-PRC1 complexes promote seed germination through H3K4me3-to-H3K27me3 chromatin state switch in repression of seed developmental genes. PLoS genetics 10: e1004091.

Møller BL. 2010. Functional diversifications of cyanogenic glucosides. Current opinion in plant biology 13: 337–346.

Narayanan NN, Ihemere U, Ellery C, Sayre RT. 2011. Overexpression of hydroxynitrile lyase in cassava roots elevates protein and free amino acids while reducing residual cyanogen levels. PloS one 6: e21996.

Narro-Diego L, López-González L, Jarillo JA, Piñeiro M. 2017. The PHD-containing protein EARLY BOLTING IN SHORT DAYS regulates seed dormancy in A rabidopsis. Plant, Cell & Environment 40:2393–2405.

Nguyen H, Schoenau JJ, Van Rees KC, Nguyen D, Qian P. 2001. Long-term nitrogen, phosphorus and potassium fertilization of cassava influences soil chemical properties in North Vietnam. Canadian journal of soil science 81: 481–488..

Nielsen LJ, Stuart P, Pičmanová M, Rasmussen S, Olsen CE, Harholt J, Møller BL, Bjarnholt N. 2016. Dhurrin metabolism in the developing grain of *Sorghum bicolor* (L.) Moench investigated by metabolite profiling and novel clustering analyses of time-resolved transcriptomic data. BMC genomics 17: 1–24.

Ogbonna AC, Braatz de Andrade LR, Rabbi IY, Mueller LA, Jorge de Oliveira E, Bauchet GJ. 2021. Large-scale genome-wide association study, using historical data, identifies conserved genetic architecture of cyanogenic glucoside content in cassava (*Manihot esculenta* Crantz) root. The Plant Journal 105: 754–770.

Panghal A, Munezero C, Sharma P, Chhikara N. 2019. Cassava toxicity, detoxification and its food applications: a review. Toxin Reviews.

Pičmanová M, Neilson EH, Motawia MS, Olsen CE, Agerbirk N, Gray CJ, Flitsch S, Meier S, Silvestro D, Jørgensen K, Sánchez-Pérez R. 2015. A recycling pathway for cyanogenic glycosides evidenced by the comparative metabolic profiling in three cyanogenic plant species. Biochemical Journal 469: 375–89.

Qian F, Zhao QY, Zhang TN, Li YL, Su YN, Li L, Sui JH, Chen S, He XJ. 2021. A histone H3K27me3 reader cooperates with a family of PHD finger-containing proteins to regulate flowering time in *Arabidopsis*. Journal of integrative plant biology 63: 787–802..

Sánchez-Pérez R, Pavan S, Mazzeo R, Moldovan C, Aiese Cigliano R, Del Cueto J, Ricciardi F, Lotti C, Ricciardi L, Dicenta F, López-Marqués RL. 2019. Mutation of a bHLH transcription factor allowed almond domestication. Science 364:1095–1098.

Santana MA, Vásquez V, Matehus J, Aldao RR. 2002. Linamarase expression in cassava cultivars with roots of low-and high-cyanide content. Plant physiology 129: 1686–1694.

Schmidt FB, Cho SK, Olsen CE, Yang SW, Møller BL, Jørgensen K. 2018. Diurnal regulation of cyanogenic glucoside biosynthesis and endogenous turnover in cassava. Plant Direct 2: e00038.

Schmidt FB, Heskes AM, Thinagaran D, Møller BL, Jørgensen K, Boughton BA. 2018. Mass spectrometry based imaging of labile glucosides in plants. Frontiers in Plant Science 9: 355425.

Shang Y, Ma Y, Zhou Y, Zhang H, Duan L, Chen H, Zeng J, Zhou Q, Wang S, Gu W, Liu M. 2014. Biosynthesis, regulation, and domestication of bitterness in cucumber. Science 346: 1084–1088.

Siritunga D, Arias-Garzon D, White W, Sayre RT. 2004. Over-expression of hydroxynitrile lyase in transgenic cassava roots accelerates cyanogenesis and food detoxification. Plant Biotechnology Journal 2: 37–43.

Siritunga D, Sayre RT. 2003. Generation of cyanogen-free transgenic cassava. Planta 217: 367–73.

Sugiyama R, Li R, Kuwahara A, Nakabayashi R, Sotta N, Mori T, Ito T, Ohkama-Ohtsu N, Fujiwara T, Saito K, Nakano RT. 2021. Retrograde sulfur flow from glucosinolates to cysteine in *Arabidopsis thaliana*. Proceedings of the National Academy of Sciences 118: e2017890118.

Szczyglowski K, Hamburger D, Kapranov P, de Bruijn FJ. 1997. Construction of a Lotus japonicus late nodulin expressed sequence tag library and identification of novel nodule-specific genes. Plant physiology 114: 1335–1346.

Wei W, Fan ZQ, Chen JY, Kuang JF, LU WJ, Shan W. 2017. A banana PHD-type transcription factor MaPHD1 represses a cell wall-degradation gene *MaXTH6* during fruit ripening. Horticultural plant journal 3:190-198.

Wei W, Huang J, Hao YJ, Zou HF, Wang HW, Zhao JY, Liu XY, Zhang WK, Ma B, Zhang JS, Chen SY. 2009. Soybean GmPHD-type transcription regulators improve stress tolerance in transgenic Arabidopsis plants. PloS one 4: e7209.

Wei W, Zhang YQ, Tao JJ, Chen HW, Li QT, Zhang WK, Ma B, Lin Q, Zhang JS, Chen SY. 2015. The Alfin-like homeodomain finger protein AL5 suppresses multiple negative factors to confer abiotic stress tolerance in Arabidopsis. The Plant Journal 81: 871–883.

Winicov I, Bastola DR. 1999. Transgenic overexpression of the transcription factor Alfin1 enhances expression of the endogenous *MsPRP2* gene in alfalfa and improves salinity tolerance of the plants. Plant physiology 120: 473–480.

Zagrobelny M, Bak S, Rasmussen AV, Jørgensen B, Naumann CM, Møller BL. 2004. Cyanogenic glucosides and plant–insect interactions. Phytochemistry 65: 293–306.

Zhang P, Bohl-Zenger S, Puonti-Kaerlas J, Potrykus I, Gruissem W. 2003. Two cassava promoters related to vascular expression and storage root formation. Planta 218:192–203.

Zhang H, Forde BG. 2000. Regulation of Arabidopsis root development by nitrate availability. Journal of experimental botany 1: 51–59.

Zhang YZ, Yuan JL, Zhang LR, Chen CX, Wang YH, Zhang GP, Li Peng, Xie SS, Jiang J, Zhu JK, Du JM, Duan CG. 2020. Coupling of H3K27me3 recognition with transcriptional repression through the BAH-PHD-CPL2 complex in *Arabidopsis*. Nature communications 11: 6212.

Zidenga T, Siritunga D, Sayre RT. 2017. Cyanogen metabolism in cassava roots: impact on protein synthesis and root development. Frontiers in Plant Science 8: 241486.

