## Supplementary material for "MePHD1.2 affects the synthesis of cyanogenic glycosides by regulating the transcription of *MeCYP79D2* in cassava": Figure & Table

**New Phytologist Supporting Information**

Article title: MePHD1.2 affects the synthesis of cyanogenic glycosides by regulating the transcriptional activity of *MeCYP79D2* in cassava

Authors: Mengtao Li[1,2], Xiao Zhao[3], Yuanchao Li[2,4], Xiaoye Zhao[5], Weitao Mai[1,2], Yajun Li[2], Qibing Liang[5], Qingchun Yin[6], Wenquan Wang[1*], Jinping Liu[1*], Xin Chen[1,2,7*].

The following supporting information is available for this article:

### Supporting Information Figures

```
>M.esculenta              v8.1|Manes.12G133500|Chromosome12:34044057..34046332
reverse|upstream=2000|downstream=0
-1505 TGTGGGAAGA TGTCCCACAT CATTAAACTA TAAAAAATTA AAAATTGTAA GTAGAAAATT
-1445 ATTAAATAAT TTAAATTAAT TATTTTAAAT TAAAGTGAGA AATGAATTTA AAAGTTATTT
-1385 AAATCAGTTT GTATTAAACT GTTTCAATTG GTATTAGAGT CATTTTGAAT GAGAAAAATT
-1325 TTATAGAGCC AGTGAACTGC GAGAAAAGAC CTATCCATGT GATACGAGGT GAAGTCGTTC
-1265 AAGTTACTGG CGAATCACAA TACTTAAGGA TTCGATCCTG ATTTAGCAAT TATATCTGAT
-1205 TCAGTTGTGA TGAGAATATC GCAAATTTAA TGGGATAAAG TTTAATAATC AGCTGCTTGA
-1145 TTAATGTGAG AACAGAATAT GTTCCACATC AGCAAAGTAT GAAAAATTAA AAATTATATA
-1085 TAAGAGTGTA ATAGTTAGCT GCTTGCTTAA TGTGGGAAGA CTTTCCACAT TAATAAAGTA
-1025 TTGAAAAATT AAAAATTATA TATAATAACT AGTATAAAAT TATTAAATAA TTTAGATTTA
 -965 ACTATTTTGA ATTAGATAAA AAGTGAATCT AAAAATTATT TGAATAAGTT TGAACCGAAC
 -905 TATTTCAATT AATCACTCAA AAATTTATAA ATAGAAAATA AAAATAGATG GATGAGGTAT
 -845 TATATATTTA AAAAATATTA AATTTATGCT ATAAAATTAA TATTTTTTAT AAATTTTAAT
 -785 ATATTGACTT ACATATAAAA TCAGTAGTTT ATTTAATTAT TTTGTCTAAT TTGATTTTTT
 -725 TAGAAAATGA TCTAATTTAA TTAACCTTAA ACTTTAATTA ATTTTACTTA ATTTTTAATA
 -665 AATAGGAATT CCTCAAAGTT GATGATAAAA GAGGGAGATT GAATCTCACA CGCCCATAAC
 -605 AAGCACTACA ACAACCTTCC CTTTCGTCTT TTTATTTTTT ATTTTTTATT TTCATTTTGC
 -545 CGTCCTTAAA GATTGAGTTT TGTAGGAGCA ATTAAATCAC GTGGCAGCCT AGAGGTTGGT
 -485 GGGCATAATG AATGAAAATC TATCTGCTTC GGCGTCAATT TTGTCAGCCA AAGTCACGTT
 -425 AAGCTAAAGG AAAACACTGC ATGGGGAGGA ATTGAAATCC TTAAAATAAA CAACGATAAA
 -365 AAAAAAATAG AAAGTTCCAA ACAGGGTCTT AATTACAGCA TTATTCTCAT TCCATCAATT
 -305 GCCTGTAAGA ATAATTCAAG AAGAAAGACC CCAGTGCCCA GAATATTATT GTATGGTTGT
 -245 CCGAAAATTA TATGAATTTA AATTATACGA GATTTTTTTT TAAATTTATT ATGTATATTT
 -185 TTCTTTAATC ATTATTCCAA TTGTTTCCCA TAATACCCTC CCCAATCCAA GGGTTAACTC
 -125 CTTAACAAAC CACTATAAAT ACCCACCCTA AGCTACGTGC ATGTTATCAA GAACAAGTCT
  -65 TCTCTTCTTC TCTTGTATGG TCTTGGTCAT AGCCCTGGAC TTGAATTGTT TAGGGCAACA
   -5 CCAAT
```

Fig. S1 Promoter sequence (2 kb) of *MeCYP79D2* gene.

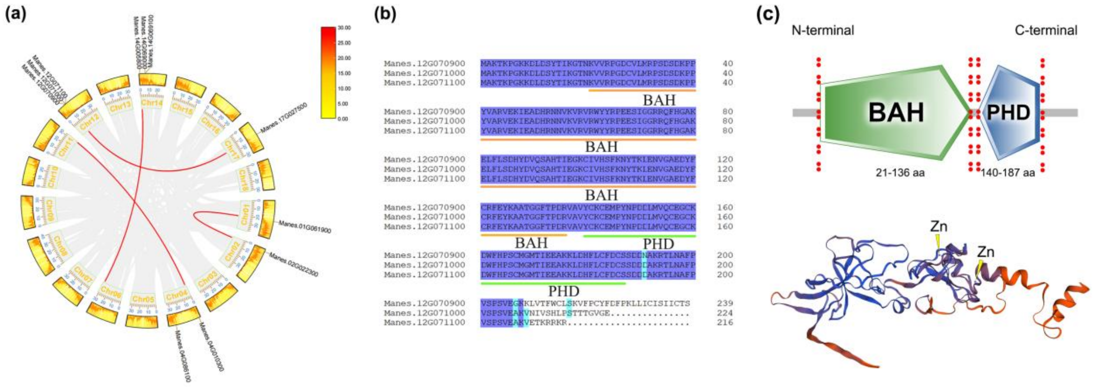


Fig. S2 Characterisation of MePHD1.2 in cassava.

(a) Phylogenetic tree of PHD family members of cassava. (b) Homologous amino acid sequence alignment in DNAMAN. (c) Conservative domain pattern diagram of MePHD1.2 protein.

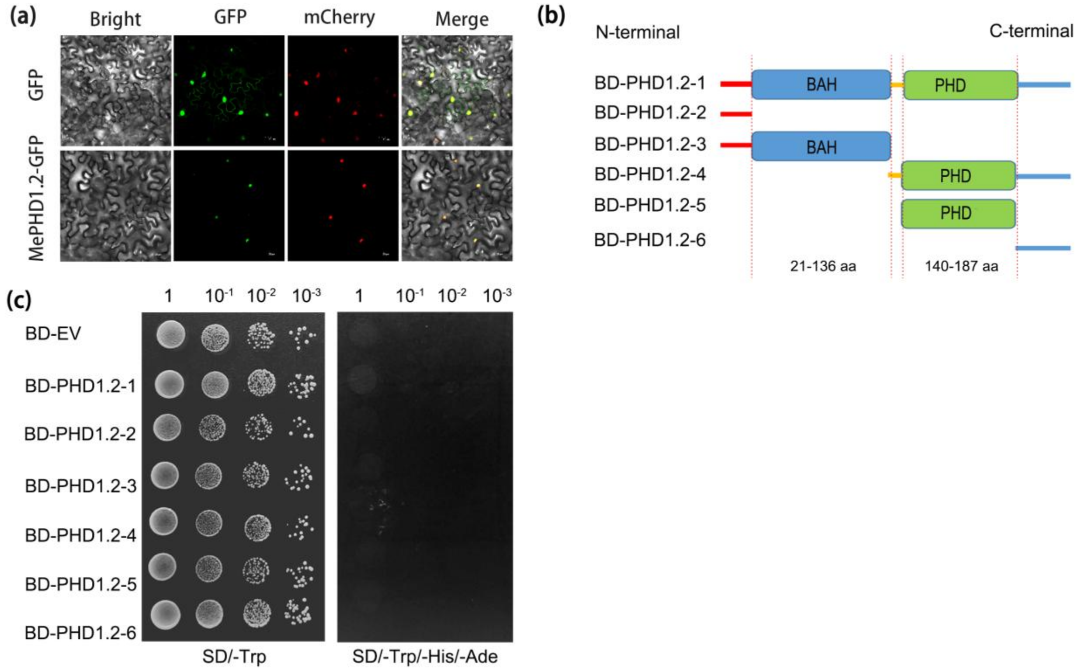


Fig. S3 Subcellular localisation and transcription activation region assay of MePHD1.2.

(a) Subcellular localization of MePHD1.2 in tobacco leaves. (b) Different truncated MePHD1.2 segments pattern diagram. (c) Verification of transcription activation activity of different truncated MePHD1.2 segments in yeast.

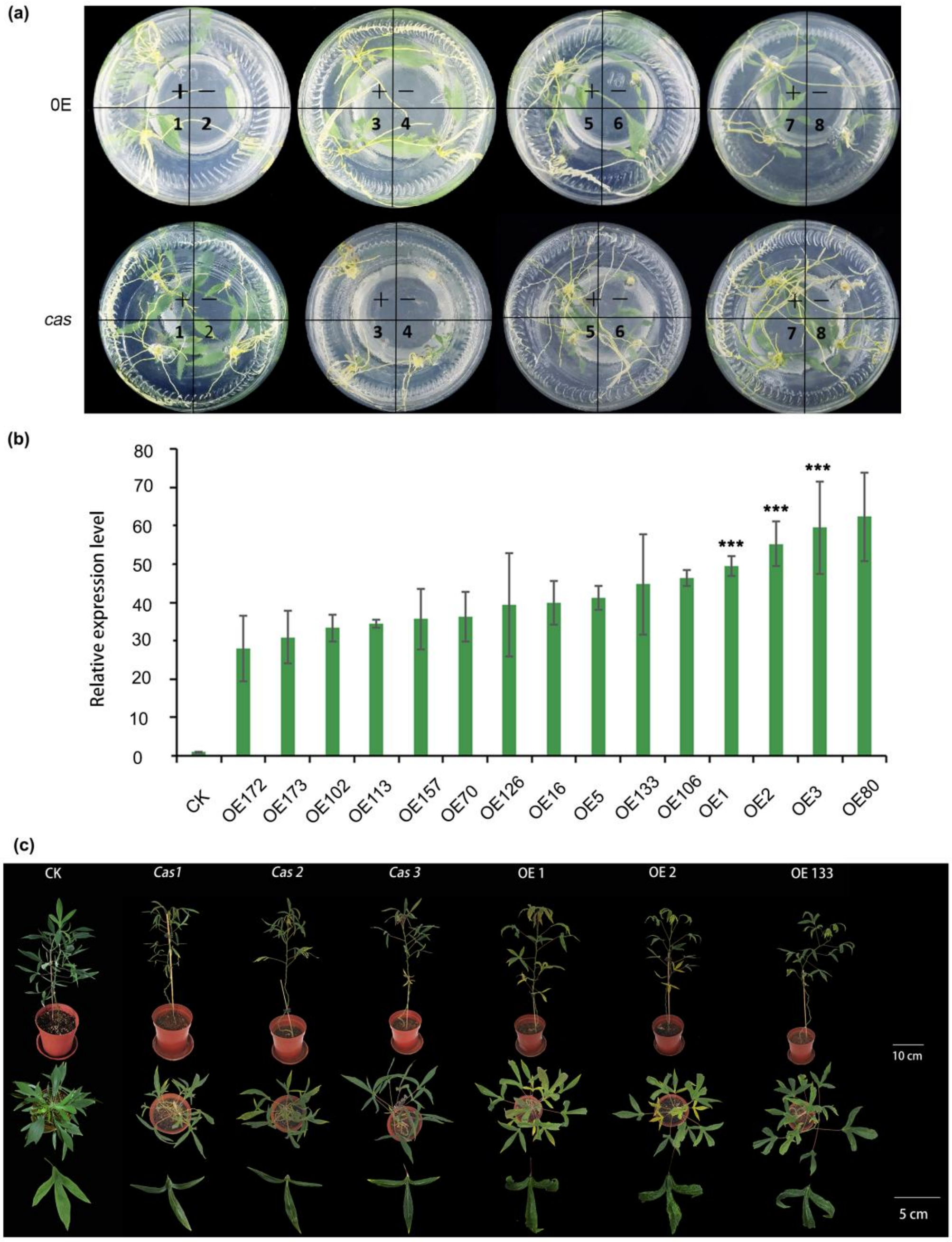


Fig. S4 Identification and phenotypic characterisation of MePHD1.2 transgenic lines and CK.

(a) Rooting screened of *MePHD1.2* tissue cultured seedlings. '+' represents Hyg resistant plants, '-' represents SC8, Hyg concentration is 20 mg/L. (b) The expression level of *MePHD1.2* overexpressed plants. (c) Phenotype of *MePHD1.2* OE and *cas* plants at 60 days after tansplanting. CK is a regenerated plant of SC8 infected with an empty vector in *Agrobacterium.*

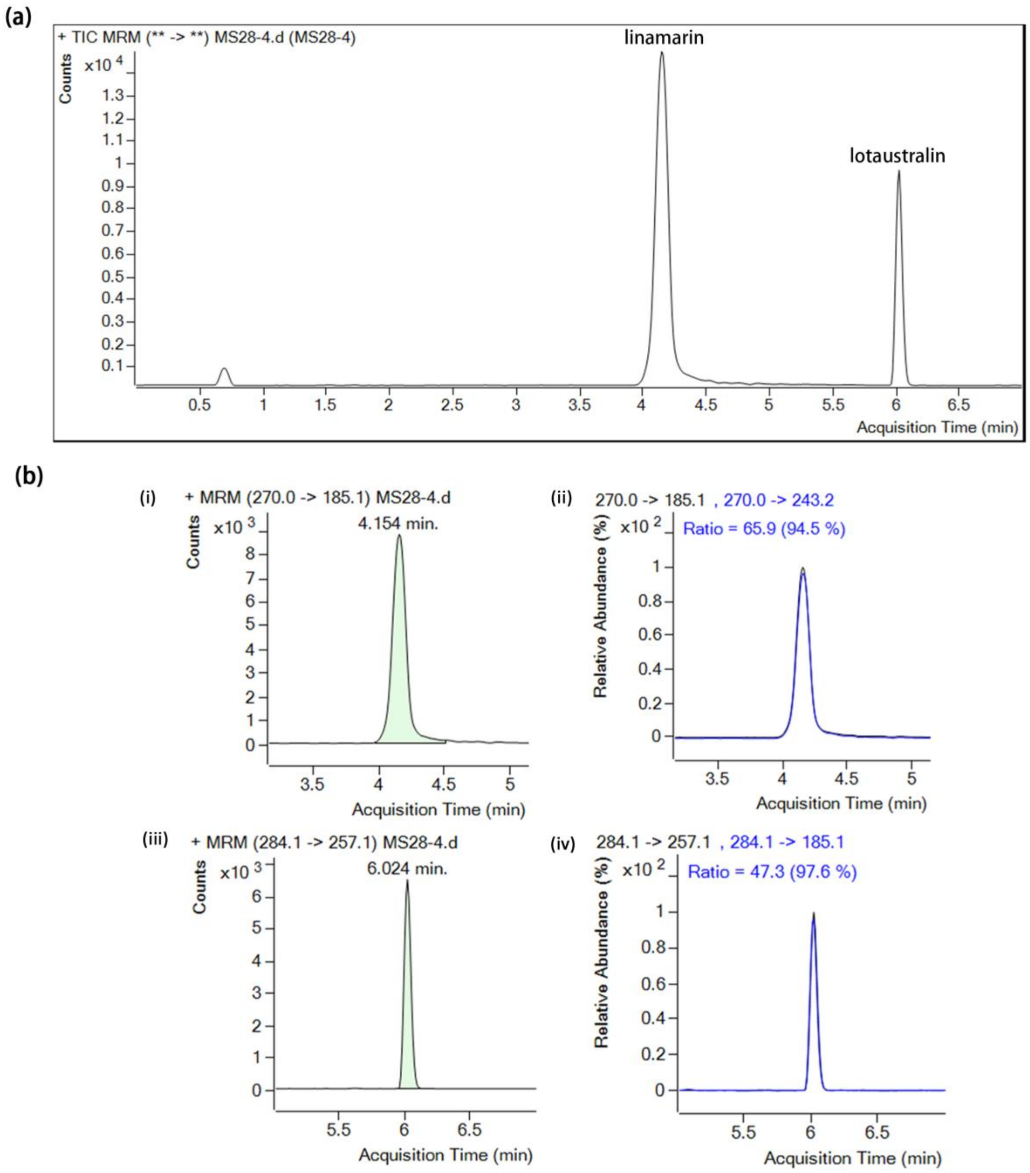


Fig. S5 MS peak of linamarin and lotaustralin determination .

(a) Mass spectrum of standard samples for linamarin and lotaustralin. (b) i-ii represents the peak time and content proportion of linamarin, respectively. iii-iv represents the peak time and content proportion of lotaustralin, respectively.

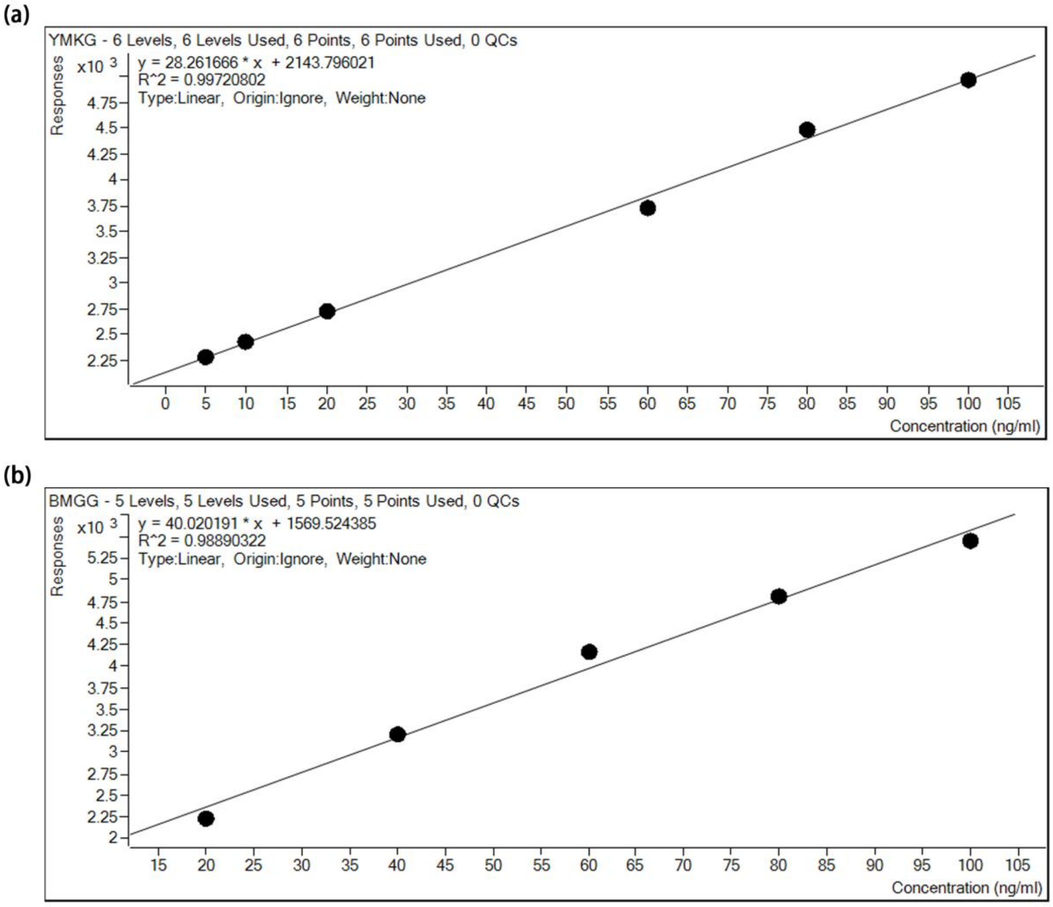


Fig. S6 Standard curves of linamarin and lotaustralin in LC-MS/MS detection

YMKG represents linamarin; BMGG represents lotaustralin.

### Supporting Information Tables

**Table S1** Part of cis-element in the MeCYP79D2 promoter.

| Component type | Sequence | Position | Length | Direction | Function Prediction |
|---|---|---|---|---|---|
| A-box | CCGTCC | -959 | 6 | + | cis-acting regulatory element |
| ABRE | ACGTG | -998, -1074, -1414 | 5 | + | cis-acting element involved in the abscisic acid responsiveness |
| ARE | AAACCA | -1386 | 6 | + | cis-acting regulatory element essential for the anaerobic induction |
| AT~TATA-box | TATATA | -414, -416, -496, -498, -660 | 6 | + | unknown |
| Box 4 | ATTAAT | -74, -359, -468, -607, -695, -959 | 6 | + | part of a conserved DNA module involved in light responsiveness |
| CAAT-box | CAAT | 877, 971, 1110, 1339 | 4 | - | common cis-acting element in promoter and enhancer regions |
| CCGTCC motif | CCGTCC | 959 | 6 | + | unknown |
| CGTCA-motif | CGTCA | 1052 | 5 | + | cis-acting regulatory element involved in the MeJA-responsiveness |
| chs-CMA1a | TTACTTAA | 823 | 8 | + | part of a light responsive element |
| G-Box | CACGTG | 997 | 6 | - | cis-acting regulatory element involved in light responsiveness |
| | CACGTT | 1074 | 6 | + | |
| LTR | CCGAAA | 1260 | 6 | + | cis-acting element involved in low-temperature responsiveness |
| TATA | TATAAAAT | 512, 689, 733 | 8 | + | unknown |
| TATC-box | TATCCCA | 330 | 7 | - | cis-acting element involved in gibberellin-responsiveness |
| TCA-element | CCATCTTTTT | 641 | 9 | - | cis-acting element involved in salicylic acid responsiveness |
| TCCC-motif | TCTCCCT | 871 | 7 | - | part of a light responsive element |
| TCT-motif | TCTTAC | 1204 | 6 | - | part of a light responsive element |
| TGACG-motif | TGACG | 1052 | 5 | - | cis-acting regulatory element involved in the MeJA-responsiveness |
| TGA-element | AACGAC | 233 | 6 | - | auxin-responsive element |

**Table S2** Primers that used for vector construction in this study.

| **Primer Name** | **Seqeunce 5' to 3'** |
|---|---|
| **Yeast one-hybrid assay** | |
| pAbA-CYP79PD2 F1 | cccaagcttTGTGGGAAGATGTCCCACAT |
| pAbA-CYP79PD2 R1 | gcgtcgacATTGGTGTTGCCCTAAACAA |
| pAbA-CYP79PD2-1 F1 | cccaagcttTGTGGGAAGATGTCCCACAT |
| pAbA-CYP79PD2-1 R1 | gcgtcgacGATTCACTTTTTATCTAATTC |
| pAbA-CYP79PD2-2 F1 | cccaagcttGCTGCTTGCTTAATGTGGGA |
| pAbA-CYP79PD2-2 R1 | gcgtcgacTATGCCCACCAACCTCTAGG |
| pAbA-CYP79PD2-3 F1 | cccaagcttGGAGCAATTAAATCACGTGG |
| pAbA-CYP79PD2-3 R1 | gcgtcgacCATATTGGTGTTGCCCTAAAC |
| pAbA-PD1 tyF1 | aaatgatgaattgaaaaagcttTGTGGGAAGATGTCCCACAT |
| pAbA-PD1 tyR1 | gcacatgcctcgaggtcgacGATTCACTTTTTATCTAATTC |
| pAbA-PD2 tyF2 | aaatgatgaattgaaaaagcttGCTGCTTGCTTAATGTGGGA |
| pAbA-PD2 tyR2 | gcacatgcctcgaggtcgacTATGCCCACCAACCTCTAGG |
| pAbA-PD3 tyF3 | aaatgatgaattgaaaaagcttGGAGCAATTAAATCACGTGG |
| pAbA-PD3 tyR3 | gcacatgcctcgaggtcgacCATATTGGTGTTGCCCTAAAC |
| pGADT7-PHD1.2 F | acgcgtcgacATGGCCAAAACCAAACCAGG |
| pGADT7-PHD1.2 R | cgcggatccTCTCTTCCTTCGCTTTGTCTC |
| pAbA F | TTCGTTCTTCCTTCTGTTCGG |
| pAbA R | ATCTCGAAAAAGGGTTTGCCA |
| 3'AD | GTGAACTTGCGGGGTTTTTCAGTATCTACGAT |
| 5'AD | CTATTCGATGATGAAGATACCCCACCAAACCC |
| **Yeast two-hybrid assay** | |
| pGBKT7-PHD1.2 F | ACGCGTCGACATGGCCAAAACCAAACCAGG |
| pGBKT7-PHD1.2 R | CGCGGATCCTCTCTTCCTTCGCTTTGTCTC |
| pGAD-PHD1.1 F | ACGCGTCGACATGGCCAAAACCAAACCAGG |
| pGAD-PHD1.1 R | CGCGGATCCTCTCTTCCTTCGCTTTGTCTCC |
| PGBKT7-F | TTGTAATACGACTCACTATAGGGCGAGC |
| PGBKT7-R | TAAGAAATTCGCCCGGAATTAGCTTGG |
| **Subcellular localization** | |
| pSL-PHD1.2 F | gcgtcgacATGGCCAAAACCAAACCAGG |
| pSL-PHD1.2 R | cgggatccTCTCTTCCTTCGCTTTGTCTC |
| **For Transcriptional Activation Assay** | |
| GAL4BD-PHD1.2 F | CGCCGTCTAGAACTAGTGGATCCATGGCCAAAACCAAACCAGG |
| GAL4BD-PHD1.2 R | TCGATAAGCTTGATATCGAATTCTCATCTCTTCCTTCGCTTTGTCTC |
| **dual-LUC** | |
| pluc-CYP79D2 F | CGGGCCCCCCCTCGAGGTCGACTGTGGGAAGATGTCCCACAT |
| pluc-CYP79D2 R | GCAGGAATTCGATATCAAGCTTATTGGTGTTGCCCTAAACAATT |

| Primer Name | Seqeunce 5’ to 3’ |
|---|---|
| 62-SK-PHD1.2 F | cccaagcttATGGCCAAAACCAAACCAGG |
| 62-SK-PHD1.2 R | cggggtaccTCATCTCTTCCTTCGCTTTGTCTC |
| **Protein Expression** | |
| C5X-PHD1.2 F | GGAATTCCATATGATGGCCAAAACCAAACCAGG |
| C5X-PHD1.2 R | ACGCGTCGACTCTCTTCCTTCGCTTTGTCTC |
| **qRT-PCR analysis for candidate genes** | |
| PHD1.2 qF | GTGCGAGGGTTGCAAAGATT |
| PHD1.2 qR | TTCACCTTGGCCTCAACAGA |
| Tubulin F | GTGGAGGAACTGGTTCTGGA |
| Tubulin R | TGCACTCATCTGCATTCTCC |
| **For Transgenic lines Construct** | |
| PHD1.2-U-F | CCCCACCTAAGCCTTCTCAATCC |
| PHD1.2-U-R | AAAGGATGGCTGGAGAAACCAA |
| pOE-PHD1-CDS F | CGCGGATCCATGGCCAAAACCAAACCAGG |
| pOE-PHD1-CDS R | ACGCGTCGACTCTCTTCCTTCGCTTTGTCTC |
| PHD-cas9 F | gattGATCTAATGGTGCAGTGCGA |
| PHD-cas9 R | aaacTCGCACTGCACCATTAGATC |
| 35S-Hyg F1 | CATTTGGAGAGGACACGCTG |
| 35S-Hyg R1 | CTATTTCTTTGCCCTCGGAC |
| Hi-PHD1.2 F | ggagtgagtacggtgtgcGGATTATTCAACTGAGACCTGGAT |
| Hi-PHD1.2 R | gagttggatgctggatggTCTTGTTAAAAATTGAGAGTCCGCA |
| **For EMSA** | |
| Biotin-M13-RVB | GAGCGGATAACAATTTCACACAGG |
| Biotin-M13-47B | GCGGTCCCAAAAGGGTCAGTGCTG |
| Biotin-PD2-1 F | GCTGCTTGCTTAATGTGGGA |
| EMSA PD2-1 R | AAATAGTTAAATCTAAATTATTT |
| Biotin-PD2-2 F | TAATTTAGATTTAACTATTTTG |
| EMSA PD2-2 R | CTATTTTTATTTTCTATTTATAAA |
| Biotin-PD2-3 F | TAAATAGAAAATAAAAATAGATG |
| EMSA PD2-3 R | CTACTGATTTTATATGTAAGTC |
| Biotin-PD2-4 F | CTTACATATAAAATCAGTAG |
| EMSA PD2-4 R | TCCTATTTATTAAAAATTAAG |
| Biotin-PD2-5 F | TTAATTTTTAATAAATAGGAATTC |
| EMSA PD2-5 R | AAAAAATAAAAAATAAAAAG |
| Biotin-PD2-6 F | CTTTTTATTTTTTATTTTTTAT |
| EMSA PD2-6 R | TATGCCCACCAACCTCTAGGC |

**Table S3** Partial candidated proteins that screening out via Y1H assay.

| Gene ID | Description | Frequency |
|---|---|---|
| **Manes.12G071000** | **PHD-finger (PHD) // BAH domain (BAH)** | **58** |
| Manes.08G021500 | TRIHELIX TRANSCRIPTION FACTOR PTL | 52 |
| Manes.18G064500 | histone H3 (H3) | 39 |
| Manes.02G076600 | HISTONE H1/H5 // SUBFAMILY NOT NAMED | 29 |
| Manes.11G137100 | histone H4 (H4) | 27 |
| Manes.15G037900 | AT-hook motif nuclear-localized protein 20 | 17 |
| Manes.18G018500 | histone H3 (H3) | 12 |
| Manes.18G088600 | pathogen-related protein | 10 |
| Manes.09G004200 | PPR repeat (PPR) | 8 |
| Manes.15G093600 | 14-3-3-like protein D | 6 |
| Manes.09G108300 | major latex allergen Hev b 5 | 6 |
| Manes.09G087400 | probable ADP-ribosylation factor GTPase-activating protein AGD6 | 6 |
| Manes.06G057800 | EIN3-binding F-box protein (EBF1_2) | 6 |
| Manes.01G201400 | DNA-directed RNA polymerase III subunit RPC8 | 6 |
| Manes.15G110900 | 60S ribosomal protein L9 | 5 |
| Manes.01G011500 | ribulose bisphosphate carboxylase small subunit, chloroplastic-like | 5 |
| Manes.15G054000 | SMALL EDRK-RICH FACTOR 1 // SUBFAMILY NOT NAMED | 4 |
| Manes.15G013200 | 60S ribosome subunit biogenesis protein NIP7 (NIP7) | 4 |
| Manes.09G072968 | syntaxin-71-like | 4 |
| Manes.07G126600 | probable aquaporin PIP-type 7a | 4 |
| Manes.17G021900 | regulator of nonsense transcripts 1 | 3 |
| Manes.13G052000 | peptidyl-prolyl cis-trans isomerase FKBP15-1 | 3 |
| Manes.11G036500 | aquaporin TIP2-1 | 3 |
| Manes.10G083600 | ADP-ribosylation factor-like protein 8 (ARL8) | 3 |
| Manes.09G068800 | aquaporin PIP2-4 | 3 |
| Manes.08G060500 | pentatricopeptide repeat-containing protein At1g80270, mitochondrial | 3 |
